# Metagenomic analyses of single phages and phage cocktails show instances of contamination with temperate phages and bacterial DNA

**DOI:** 10.1101/2024.09.12.612727

**Authors:** Xue Peng, Sophie Elizabeth Smith, Wanqi Huang, Jinlong Ru, Mohammadali Khan Mirzaei, Li Deng

**Affiliations:** Technical University of Munich, Germany; TUM School of Life Sciences, Professor for Prevention of Microbial Diseases; Central Institute of Infection Prevention (ZIP), Freising, 85354, Germany; Institute of Virology, Helmholtz Centre Munich - German Research Centre for Environmental Health, Neuherberg, 85764, Germany

## Abstract

Increasing antibiotic resistance has led to renewed attention being paid to bacteriophage therapy. Commercial phage cocktails are available but often their contents of the phages are not well defined. Some metagenomic studies have been done to retrospectively characterise these cocktails, but little is known about the replication cycle of the included phages, or about the possible bacterial DNA contamination. In this study, published metagenomic sequences were reanalysed using recent advances in viromics tools. Signs of temperate phage contigs were found in all cocktail metagenomes, as well as host DNA, which could poses a risk as it may lead to horizontal gene transfer of virulence factors to commensals and pathogens. This suggests the need to implement further quality measures before using phage cocktails therapeutically.

## Introduction

Antibiotics were once lauded as the end of bacterial diseases, however, in the face of ever more multi-drug resistant pathogens and a dearth of new antibiotics in the drug discovery pipeline, it is becoming increasingly clear that we are approaching a post antibiotic era. In 2019, it is estimated that more than a million deaths were directly caused by Antimicrobial Resistance (AMR) infections (1) and this number is expected to rise to 10 million by 2050 (2). Due to the urgent need for alternative methods of treating multi-drug resistant infections, much interest has turned to the use of bacteriophages, or phages, for this purpose. In the west, phage therapy largely remains at a pre-clinical stage and is only authorised for compassionate use when all other options have failed. Elsewhere, the Eliava Institute in Tbilisi, Georgia, leads the way in providing commercial phage products for the treatment of infections that have failed to be treated by more conventional means (3).

There are a number of ways in which phages differ from traditional antibiotics, including that they are highly specific, often only able to infect one or two bacterial strains (4). This can be a problem when treating a bacterial infection caused by an unknown host. There are two possible approaches to combat this – the first is to culture the infecting bacteria and test potential phage treatments to ensure susceptibility (5). However, due to the time critical nature of advanced and serious bacterial infections, this approach can be impractical. Instead, a commonly used approach is to administer a cocktail of phages which, together, are able to infect a broad range of potential hosts (6). Phage cocktails might also have the advantage of hindering the development of resistance, as bacteria resistant to one phage will likely be killed by another in the cocktail (6). As such, much consideration should be given to intelligent cocktail design, so that a final cocktail of well characterised phages is as efficient, broad host range, and resilient against resistance development as possible (6).

In addition, obligately lytic phages are preferred for phage therapy, and temperate phages, which are able to enter the lysogenic cycle and incorporate themselves into the bacterial genome, are not considered. This is due to concerns that they have the ability to facilitate horizontal gene transfer and may provide virulence factors such as toxins or AMR genes to the host (7). Several bacterial diseases including diphtheria, cholera and dysentery encode the toxin in question on a prophage (8). Prophages may also interact with other phages, including those used for phage therapy, protecting their hosts from infection with other phages (9). It is reasonably easy to select only virulent phages for use in phage therapy, as phages with genes allowing them to enter the lysogenic cycle, such as integrases and excisionases, can be identified bioinformatically from the phage genome (10–12). However, this information is not always available for the commercial cocktails available in the market. Generally, commercialy available products are wide ranging cocktails based on the suspected pathogen or site of infection and these have been used to treat patients for many years (3, 13, 14). Some phage treatments are also available under compassionate use guidelines (15). Despite this, it is unknown exactly which phages are contained in some commercial cocktails, and how they were characterised or selected for use. In addition, sequences are often not publicly available, and little is known about their replication cycles.

Moreover, as phages require a bacterial host in order to replicate, it is possible for phage cocktail preparations to become contaminated with temperate phages induced from the propagating host. Prophages are induced spontaneously at a rate of approximately one in 10^2^ - 10^4^ cells (16), and further prophage induction can be caused by any conditions which would favour the lytic replication cycle, for example, high host density (17), or stressors such as UV light (18). Treatment with virulent phages has also been known to induce prophages from the host (18), and this poses a problem for the process of preparing phages for phage therapy. It is common for phage preparations to contain impurities and bacterial debris by virtue of the process required to grow the included phages, for example they often contain endotoxins released by lysed bacteria and must be purified before use (19).

It has long been accepted that temperate phages should not be used for phage therapy, but until recently little attention has been paid to the risks posed by application of these phages. However, there is some evidence that the presence of temperate phages in bacteriophage cocktail preparations is not as serious an issue as it might seem, and may not have a direct negative impact on a cocktail’s efficacy, although evidence is mixed on this topic (18, 20). To remedy the lack of available information about commercial cocktails, several papers have attempted to retrospectively characterise them through metagenomic analysis (5, 21–23). To explore whether temperate phages can be found in phage cocktails, we have revisited published sequences from commercial phage cocktails and reanalysed them using recent advances in viromics approaches and tools. We have also investigated individual phages we have sequenced ourselves for evidence that phage enrichment might lead to the induction of prophages from the bacterial host that was used for propagation.

## Results

### Commercial phage cocktails

Sequences of four cocktails available commercially from the Eliava Institute were sequenced previously. Three were PYO cocktails (PYO97, PYO2000 and PYO 2014), which purports to contain phages against a wide range of pathogens and is intended for the treatment of burn wound, respiratory, gastrointestinal and other infections. The fourth is an INTESTI cocktail, which is for the treatment of gastrointestinal infections. The metagenomic data from the cocktails were assessed to determine whether they contained temperate phages (Table 1) based on a range of measures, including the presence of genes associated with a lysogenic lifecycle such as integrases and excisionases. A total of 89 contigs were classified as temperate, of which 6 contained an integrase gene – 2 in the PYO97 cocktail and 4 in the PYO2014 cocktail (Tables 1, S3 and S6 and Figure S4 B). According to CheckV analysis, none of these sequences contain host genes, suggesting they are not contaminating host DNA. However, to achieve absolute certainty, it is crucial to obtain the complete genome. The relative abundance of these sequences categorised as temperate phages is very low (Table S3). A comparative genomic analysis was performed to compare each of these sequences with its closest match in NT database (https://academic.oup.com/nar/article/42/D1/D7/1054454) identified using BLASTn and amino acid level comparisons were done by Clinker (Figure 1).

**Table 1.** Relative abundance (%) of temperate and virulent sequences in the four commercial cocktails, After relative abundance values, the absolute number of contigs is shown in brackets.

|  | PYO97 | PYO2000 | PYO2014 | INTESTI | 123 |
| --- | --- | --- | --- | --- | --- |
| **Accession number** | ERR2184199 | ERR2184200 | ERR2184201 | SRR3744915 |  |
| **mean raw reads length** | 243 | 248 | 244 | 177 | 124 |
| **Total raw reads base** | 2 333 659 551 | 629 863 716 | 8 673 605 455 | 166 115 639 |  |
| **Total number of raw reads** | 9 575 268 | 2 533 708 | 35 454 908 | 938 802 | 125 |
| **Total clean reads base** | 2 057 343 861 | 249 631 025 | 7 458 026 778 | 157 824 205 |  |
| **Total number of clean reads** | 8 492 056 | 1 037 354 | 30 694 542 | 863 326 |  |
| **Total number of contigs (≥1kb)** | 470 | 247 | 755 | 225 | 126 |
| **Min length (bp)** | 1000 | 1005 | 1000 | 1003 |  |
| **Max length (bp)** | 344 583 | 93 033 | 212 402 | 128 699 | 127 |
| **Number of binned sequences** | 133 | 55 | 256 | 82 |  |
| **Number of bins** | 35 | 13 | 46 | 23 |  |
| **Total number of viral contigs** | 256 | 152 | 380 | 196 | 128 |
| **Number of viral contigs above medium quality** | 19 | 6 | 16 | 15 |  |
| **Average length of viral contigs above medium quality (bp)** | 8 5917.32 | 53 214.17 | 73 865.00 | 67 443.13 | 129 |
| **Maximum length of viral contigs above medium quality (bp)** | 344 583 | 93 033 | 212 402 | 128 699 | 130 |
| **Median length of viral contigs above medium quality (bp)** | 47 144 | 54 539 | 47 897 | 58 371 |  |
| **Minimum length of viral contigs above medium quality (bp)** | 5441 | 5485 | 38 562 | 28 300 | 131 |
| **Relative abundance of temperate viral contigs** | 0.1674 (31) | 0.0949 (4) | 0.6998 (45) | 1.8015 (9) |  |
| **Relative abundance of temperate viral contigs with integrase gene** | 0.0173 (2) | - | 0.3926 (4) | - | 132 |

**Figure 1.**
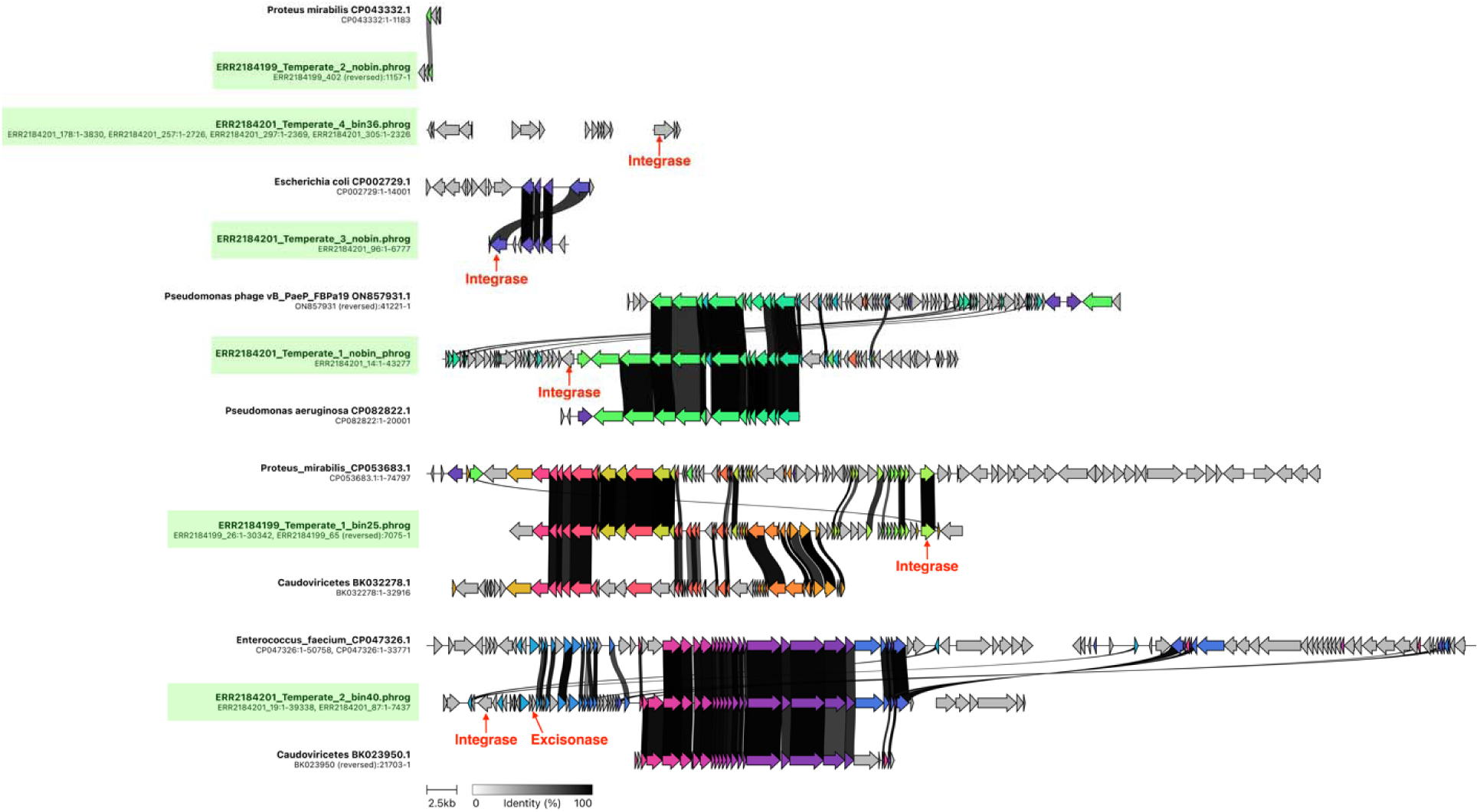
Comparative genomic map of four temperate contigs identified in cocktail sequences with the most similar sequences found in the NT database. The minimum identity of connections between genes is 0.6. Lysogeny-related genes are labeled.

We also tested whether there was bacterial contamination in the phage cocktail sequences. Some bacterial fragments could be found in the PYO cocktail sequences, mostly from *Escherichia coli* and *Shigella flexneri*, whilst the INTESTI cocktail contained very little bacterial contamination, showing that there is inconsistency in the amount of contamination between different cocktails. The bacterial reads could be mapped to the entire length of the bacterial genome, that is, the sequences are not exclusively from the prophage regions of the reference genomes, and are therefore more likely to come from contaminating DNA than prophages. However, in these cocktails the number of bacterial reads remaining after quality filtering is very low (Table S2).

### Individually sequenced phages

Given the lack of knowledge of how the phages in the commercial cocktails were isolated and characterised, individual phages which had been isolated using standardised protocols were sequenced and analysed. These phages were treated with DNAse during the DNA extraction process to reduce host contamination. Different bacterial species were selected for phage isolation and propagation to evaluate the role of the host bacteria in contamination, and unlike for the commercial cocktails, the hosts were known and could be used for further analysis. Phages were initially characterised *in vitro* in terms of plaque morphology, host range, and morphology (Figure 2B, Figure S3) before being sequenced individually for genomic characterisation (Figure 2A, 2C, and S4A). HMGUsm2 contained temperate phage contigs, but at a low abundance (Table 2), alongside a virulent genome at a higher abundance, which was therefore likely to be the target phage.

**Figure 2.**
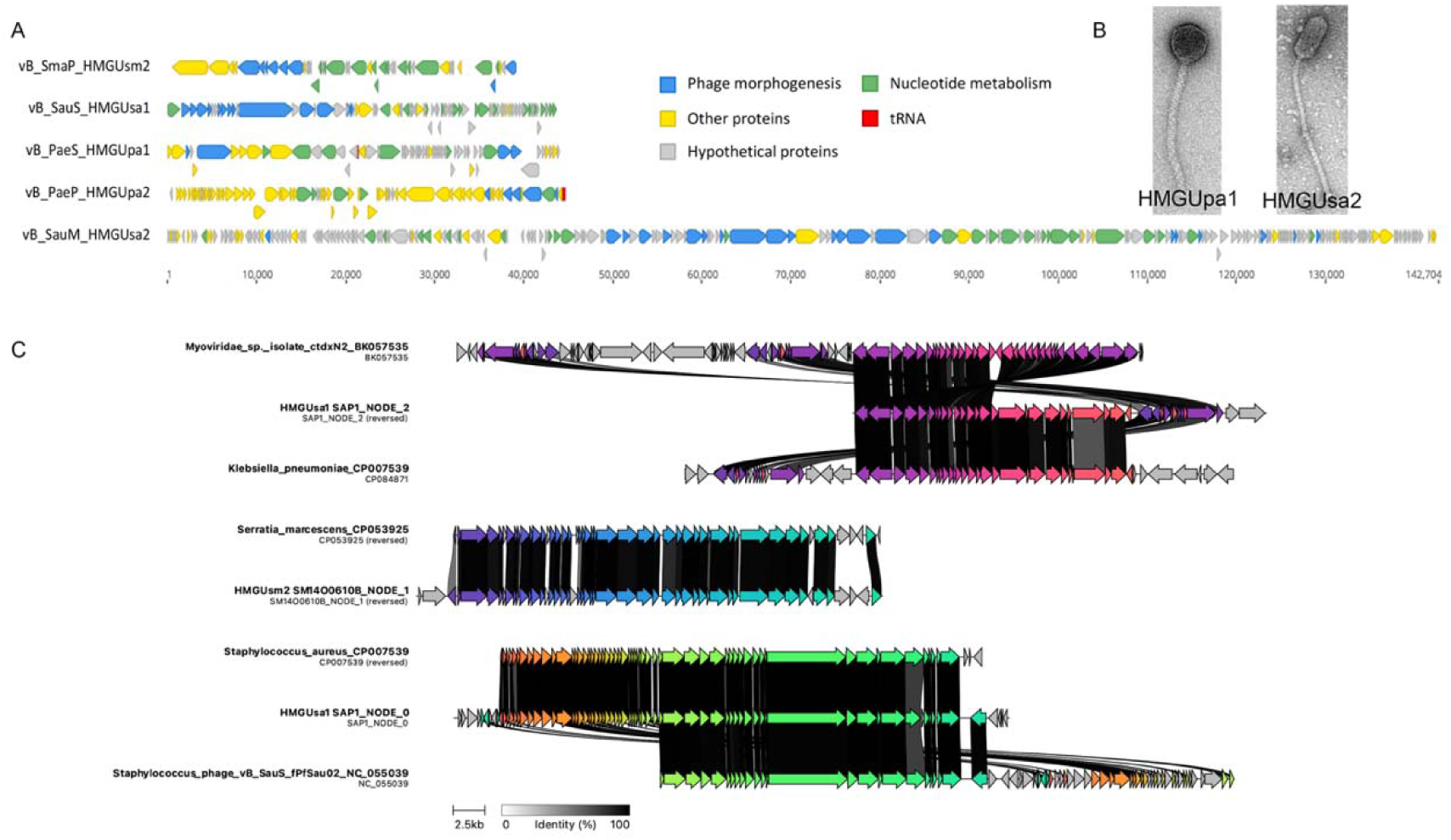
A) Genomic map of the individually sequenced phages. Genes associated with phage morphogenesis are coloured blue, those associated with nucleotide metabolism are coloured green, tRNA genes are coloured red, and other known proteins are coloured yellow. Unknown hypothetical proteins are coloured grey. B) TEM images of HMGUpa1 and HMGUsa2. C) Comparative genomic map of temperate contigs identified in individually sequenced phage sequences with the most similar sequences found in the NCBI database. The minimum identity of connections between genes is 0.6.

**Table 2.**
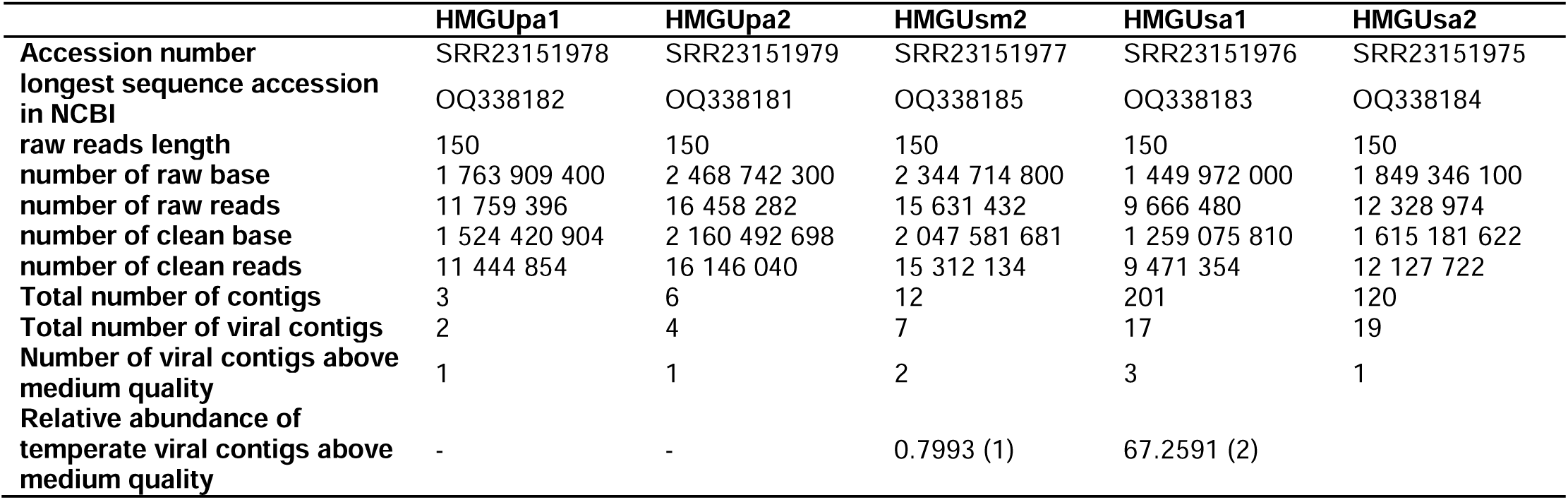
Relative abundance (%) of temperate and virulent sequences in the individually sequenced phages. After relative abundance values, the absolute number of contigs is shown in brackets.

HMGUsa1, on the other hand, contained two integrase-gene-containing temperate contigs, and it was a temperate contig that was at a high abundance (67.26%). This suggests that the sequenced phage is indeed temperate, despite experimental indicators to the contrary, such as non-turbid plaques (Figure S3A), demonstrating the need to sequence phages before use. HMGUsa2, HMGUpa1 and HMGUpa2 only contained lytic phage genomes. Contigs containing the endolysin gene were reconfirmed as temperate by comparing them to their most similar match in the NT database, including both bacteria and phage, using BLASTn (Figure 2C, Table S3).

The individually sequenced phages were treated with DNase during the DNA extraction process, a standard and common step used to reduce the amount of host DNA contamination. To discover whether this treatment was successful in removing DNA contamination, the sequences were analysed for the presence of bacterial contamination. Where bacterial contamination was present, it was mostly of the host bacteria, as might be expected. However, HMGUsa1, an *S. aureus* phage, also contained *Serratia marcescens* contamination (Figure S2). When the sequences were analysed for the presence of virulence factors, six virulence genes were identified in bacterial contigs present in HMGUsa2 (Table S1). To more effectively compare the results from the individually sequenced phages to the commercial cocktails, a cocktail was simulated *in silico*. After assembly, 7 contigs were identified which were medium quality or above by CheckV. Of these, 4 were classified as virulent, and the remaining 3 were classified as temperate. This concurs with the assessment of the individual sequences, where 4 of the phages were classified as virulent, 1 as temperate, and an additional 2 temperate contigs were identified (Table S5).

## Discussion

In both of the commercial cocktails, as well as some of our own sequenced virulent phages, temperate phage contigs were present, and similar results have been found in previous metagenomic studies of commercial cocktails (22). Since it is widely accepted that temperate phages are not suitable for phage therapy, and considerable efforts are made to exclude them from phage cocktails, it seems unlikely that the temperate phages were deliberately included in the commercial cocktails. Therefore, there are three possible sources for the temperate phage genetic material: (1) the phages were wrongly identified as virulent phages; (2) there were free temperate phages in the cocktail, likely induced from the propagating host during the preparation of individual phages before they are combined into a cocktail; or, alternatively, (3) the temperate phage sequences were from contaminating host DNA.

The first possibility is that a temperate phage was misidentified as virulent and subsequently included in the cocktail. The individually sequenced phages, and presumably the phages included in the commercial cocktail as well, were isolated through classical approaches and were initially characterised using plaque morphology and kinetics. Many temperate phages can be identified with these methods, as they would be expected to have turbid plaques and inefficient bactericidal abilities. However, this is not universally true, as can be seen by the presence of lysogeny related genes in HMGUsa1, a phage which had been experimentally identified as virulent. In the absence of this genomic information, this phage would have been considered for therapeutic use. Not all commercially available phage cocktails are sequenced (15, 24–26), which may pose risks. HMGUsa1 is not the first example of a temperate phage forming clear plaques (27) and although it is unclear how common this is, this result demonstrates the importance of including a sequencing step when characterising a phage for use in phage therapy. This step is required by the Belgian Institute for Public Health (Sciensano), which leads the way in developing a pipeline for phage therapy approval in Europe, in order to issue a ‘green genomic passport’ and certify a phage safe for therapeutic use (28).

A second possible explanation for the presence of temperate phage contigs is that they were present as a result of host DNA contamination in the phage lysate. Although the level of host DNA contamination in the cocktails is low, it can be seen in most of the individually sequenced phages. Many bacteria, including pathogens such as *Streptococcus pneumoniae*, *Vibrio cholerae*, and *Haemophilus influenzae* (29), are naturally competent and are able to take up free DNA in their environment. This means that host DNA that is present in the lysate of a phage used for phage therapy could potentially be taken up and used by the host microbiota. Yet, little is known about the effect of bacterial DNA in phage cocktails, and there is no definition of how low the host DNA level should be in order to not pose a problem. Although host DNA contamination may explain the presence of some temperate phage contigs, at least 6 of the contigs contain prophage related genes and may instead come from induced prophages, or at least from prophages in contaminating host DNA. Prophages are often induced in response to a stressor, such as exposure to antibiotics or phage predation. In addition, spontaneous prophage induction happens at a low but constant rate, so even if the virulent phage treatment itself did not directly induce prophages, it is possible that phage preparations would always contain free temperate phages if the host were a lysogen. The only sure way to avoid this would be to use a host from which prophages could not be induced: this could be either because it was free from prophages, or that it only contained cryptic prophages, be they naturally occurring or engineered. However, given the high prevalence of prophages in pathogens, and the differences in infection efficiency between different hosts, finding such a host which is also efficient at producing high titres of the phage can be challenging. However, there are some alternatives. If the therapeutic phage has a broad host range, it may be possible to use a non-pathogenic propagating host which would be less likely to contain prophages. Alternatively, a cell-free system could be used to manufacture phages using only the extracted replication machinery from the bacterial host. Although cell-free systems are not currently a realistic way to mass produce phages, as they have only been used effectively on a limited number of model phages, and face challenges reaching high enough concentrations (30), this approach holds much potential. Until this tool becomes economically viable, all potential hosts should be sequenced and certified that they contain no inducible prophages and are safe to use for propagation of therapeutic phages.

The relative abundance of the sequenced temperate phages is very low in both the cocktails and the individually sequenced virulent phages (Table 1), which suggests that even if prophages are being induced, they remain at a very low concentration in comparison to their virulent counterparts. It has been suggested that this may be due to difficulties in replicating when they must compete for hosts against the more abundant virulent phages, and that at such a low concentration, they are unlikely to have a negative impact on treatment outcomes. Indeed, the PYO cocktail has, amongst others, been used effectively for a number of decades without raising any significant safety concerns (3). Despite suggestions that the presence of temperate phages in treatments may lead to horizontal gene transfer and drive the development of more virulent, more extensively drug resistant pathogens, there seems to be little evidence of this in practice. Recently, the PYO cocktail was used together with antibiotics to successfully treat a cystic fibrosis patient with a persistent *P. aeruginosa* infection. In this case, the bacteria in question were observed to become less virulent, with an increased susceptibility to some antibiotics (3). There are also many other positive examples of phages being used to successfully treat patients. However, many of them are case studies of individual patients rather than full scale studies (3, 13). There is very little information on the long-term effect of these cocktails, and on whether evidence of horizontal gene transfer might be apparent on a longer timescale.

The possibility of inducing prophages in the propagating host may prove hard to rule out, but could be solved by using a host without prophages. Licencing authorities may consider certifying sequenced phages and their hosts to ensure they are safe for therapeutic use, as is done by the Belgian Institute for Public Health. They could also consider defining a limit for an acceptable concentration of both unintended induced prophages and bacterial DNA in a phage cocktail. In industrial settings, standard procedures for protein production include using a bacterial strain with prophages knocked out, and analysing the sample at multiple stages for the presence of phages: perhaps a similar process would also be of benefit in the production of therapeutic phages. Although a sequencing step is now common practice when characterising phages intended for therapeutic use, it rarely assesses their level of contamination with prophages and bacterial DNA. Therefore, we would suggest that all phage cocktails be fully characterised and evaluated for potential contamination before therapeutic use, and that detailed information on their contents, including sequencing data, should be publicly available. This problem is not unusual, and even well-defined cocktails produced in the west have not previously considered the potential risk contaminating prophages or host DNA might pose.

## Materials and Methods

### Bacterial strains and bacteriophages

The bacterial strains used as hosts in this study can be found in Table 3. *Pseudomonas aeruginosa* strain 50071 was purchased from the DSMZ, *Serratia marcescens* strain NRZ-49541 was provided by Dr. Jörg B. Hans (RUHR University, Germany), and *Staphylococcus aureus* strains SH1000 and Newman were provided by Prof. Andreas Peschel (University of Tübingen, Germany). Additional strains used for host range testing can be found in Table S4. All bacteria and phages were cultivated in LB broth (Luria/Miller, Carl Roth, Germany) at 37 °C with continuous shaking at 200 rpm. In total, there were five phages isolated and sequenced individually: *P. aeruginosa* phages HMGUpa1 and HMGUpa2; *S. marcescens* phage HMGUsm2; and *S. aureus* phages HMGUsa1 and HMGUsa2.

**Table 3.** Bacteriophages and their bacterial hosts used in this study.

| Bacterial Strain | Species | Source | Phages infecting this host |
| --- | --- | --- | --- |
| DSM 50071 | *Pseudomonas aeruginosa* | DSMZ | HMGUpa1 |
|  |  |  | HMGUpa2 |
| NRZ-49541 | *Serratia marcescens* | Dr. Jörg B. Hans (RUHR University, Germany) | HMGUsm2 |
| SH100 | *Staphylococcus aureus* | Prof. Andreas Peschel (University of Tübingen, Germany) | HMGUsa1 |
| Newman |  |  | HMGUsa2 |

### Phage isolation, propagation and concentration

All phages in this study were isolated from wastewater according to the following procedure: 50 ml of wastewater was filtered through a 0.45 µm pore filter (Merck, Germany) to remove bacteria. The filtrate was mixed with an equal volume of 2 x LB broth and inoculated with 1% bacterial overnight culture. The enrichment culture was incubated at 37 °C overnight shaking at 200 rpm. Subsequently, the culture was centrifuged for 30 minutes at 4 500 x g and filtered. The presence of phages was assessed by plaque assay as described previously (31). Each phage was purified by four rounds of plaque purification using plaque assays.

Each phage was propagated on the appropriate host strain in LB broth at 37 °C. Briefly, 300 mL of LB broth was inoculated with 5 mL bacterial overnight culture and incubated for ca. 3 hours until the bacteria reached exponential phase. Then, the bacterial culture was infected by 2 mL of phage lysate in LB and incubated overnight. Subsequently, the enrichment culture was centrifuged at 4 000 x g for 20 minutes and the supernatant was transferred into High-Speed centrifuge tubes (Thermo Scientific^TM^, USA). The propagated phages intended for sequencing were 1:100 concentrated by centrifugation at 35 000 x g for 2 hours at 4 °C, using an Avanti J-E High-Speed Centrifuge (Beckman, USA). The pellet was resuspended in 3 mL SM buffer (0.1 M NaCl, 10 mM MgSO4•7H2O, 50 mM Tris-HCl, pH 7.5) and filtered through a 0.22 µm filter. The phage titre was determined by plaque assay.

### Extraction of phage DNA and phage genome sequencing

Phage DNA was extracted from the concentrated phage stocks using the modified phenol-chloroform method. In brief, 600 µL of phage sample was treated with 2 µL (4 units) of TURBO^TM^ DNase (Thermo Fisher, USA) at 37 °C for 1.5 hours to remove external DNA. DNAse was inactivated by incubation at 65 °C for 30 minutes. The phage capsid was digested by 40 µL of 20 mg/mL proteinase K (Thermo Fisher, USA) and incubated for 1 hour at 37 °C. Subsequently, the sample was mixed 1:1 with phenol:chloroform:isoamyl alcohol (25:24:1, v/v) (Sigma-Aldrich, USA) and phage DNA was separated from proteins in 5PRIME Phase Lock Gel-Light tubes (QuantaBio, USA) by centrifugation at 12 000 x g for 5 minutes. DNA in the aqueous phase was washed in 100 % ice cold ethanol and incubated at -20 °C overnight. Phage DNA was precipitated after centrifugation at 18 000 x g at 4 °C for 1 hour. After removing the ethanol, the DNA pellet was dissolved in TE buffer. The extracted phage DNA was purified using Genomic DNA Clean & Concentrator-25 kit (ZYMO Research, USA). DNA concentration was determined by Qubit^TM^ dsDNA HS Assay Kit (Thermo Fisher, USA). Genomic DNA of the 5 phages was sequenced separately by Novogene. The NEBNext® DNA Library Prep Kit (New England BioLabs, US) was used to prepare the samples for sequencing using the Illumina platform (NovaSeq 6000, PE150).

### In vitro phage characterisation

#### Transmission Electron Microscopy (TEM)

Phage lysates were concentrated to at least 10^10^ PFU/mL by centrifuging at 35,000 x g for 3 hours before being resuspended in SM buffer. 5 µL of the sample was blotted onto a carbon-coated grid. After 1 minute of incubation, the grid was stained with 5 µL of 2% uranyl acetate for 30 seconds. The phages were visualised by a JEOL JEM-1400 Plus transmission electron microscope operated at 80 kV and 50 000 x magnification.

#### Host range

Host range was assessed using spot tests. 100 μL of bacterial culture was added to 3 mL of 0.7% agar and spread over an LB plate to form a bacterial lawn. Phage lysate was serially diluted and 10 μL of each dilution was spotted onto the top of the bacterial lawn. Phages were considered to infect the host only if individual plaques could be seen.

### Cocktail metagenomes from public databases

Public data from four previously analysed cocktails (PYO97, PYO2000, PYO2014, INTESTI) were downloaded from NCBI (ERR2184199, ERR2184200, ERR2184201, SRR3744915) (32). These cocktails had been obtained from the Eliava Institute in Georgia and sequenced as described previously (5, 23) – their published raw reads were used for this study.

### Bioinformatic Analysis

All of the following steps were performed on both the Eliava cocktails and the individual sequenced phages unless otherwise specified. Code can be found at https://github.com/deng-lab/ProphageCocktail

### Quality Control and Assembly

A PhiX sequencing control had been used during the sequencing of all of the PYO cocktails, and PhiX sequences could be identified in PYO97 (8.43%), PYO2000 (23.07%), and PYO2014 (2.81%). PhiX sequences were identified and removed from raw reads using Bowtie2 (v2.3.5.1) (33). Then, Fastp (v0.23.2) (34) was used to control the read quality, before the clean reads were assembled using SPAdes (--meta, v3.15.2) (35). Contigs with a length shorter than 1000 bp were removed. Since VirSorter2 (36) tends to overestimate prophage size, CheckV (v0.8.1) (37) was first used to remove host DNA and assess the quality of the contigs, before VirSorter2 was used to identify the viral contigs after the host region had been removed. For the cocktail sequences, 2 criteria were applied to make sure the assembled contig is a viral contig: 1) the contig quality should not be equal to “not-determined” as assessed by CheckV, and 2) the contig should be classified as a virus by VirSorter2. Contigs which met these two criteria were used for future characterisation analysis. For the individually sequenced phages, the following alternative criteria were used: 1) contigs should be above medium quality as assessed by CheckV, and 2) contigs must be classified as viruses by VirSorter2. The discrepancy in criteria between the two sets of data are due to the simpler composition of the individually isolated and sequenced phages compared to the cocktails. We also used vRhyme (v1.1.0) (38) with the default settings to bin contigs longer than 1000 bp after assembly. Assembled phage contigs were annotated using RAST (https://rast.nmpdr.org/).

### Relative Abundance

In order to have a more precise abundance estimation, the clean reads were mapped to contigs after CheckV using Bowtie2 (v2.3.5.1) (33). The number of mapped reads were calculated using SAMtools (v1.13) (39). The relative abundance was calculated using the following formula:

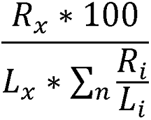

in which R_X_ denotes the number of reads mapped to a contig, L_X_ is the length of the contig, and) 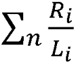 corresponds to the sum of mapped reads (R_L_) normalised by contig length (L_L_ ).

Comparative Genomic Analysis Replidec was used to predict the replication cycle of phages. And for 6 temperate phages contigs with integrase genes were chosen, to manually align to the NT database using webversion BLASTn (default parameter) (Supplementary Table S3) (40) and And the bacteria genomes or viral genome in NT database with the highest alignment score and identity greater than 95% were downloaded. Also Clinker (v0.0.23) (41) was applied to check the amino acid level similarity with a loose parameter (-i 0.3) to ensure no remote homologs. In the Clinker figure, similarly above 0.6 were connected with line between genes.

### Assessment of bacterial contamination

To assess if there was bacterial contamination of the sequences, reads that could not be mapped to the CheckV and VirSorter2 results were recruited. Then, these reads were classified using Kraken2 (v2.1.2) (42) with the MinusB (v 12_9_2022) database. The bacterial genomes with the most reads mapped to them were retrieved from NCBI and Bowtie2 (v2.3.5.1) was used to confirm that the reads mapped randomly across the length of the genome. Genomes of *Enterococcus faecium* (GCF_009734005.1), *Escherichia coli* (GCF_000005845.2, GCF_000008865.2), *Proteus mirabilis* (GCF_000069965.1), *Shigella flexneri* (GCF_000006925.2), *Pseudomonas aeruginosa* (GCF_000006765.1), *Serratia marcescens* (GCF_003516165.1), and *Staphylococcus aureus* (GCF_000013425.1) were downloaded and analysed.

### Simulation of phage cocktail *In Silico*

Firstly, used the clean reads of the 5 individually sequenced phages were normalised separately using reformat.sh and bbnorm.sh (target=100, min=5) in BBTools (v38.18) (56). After merging the normalised clean reads, the same assembly procedure as with the commercial cocktails was followed: first SPAdes (--meta, v3.15.2) was used for assembly, before CheckV (v0.8.1) was used to remove host DNA and assess the quality of the contigs. After this, Replidec (v0.2.3.1) was used to predict the replication cycle of assembled contigs to determine if temperate phages could be detected in the mock phage cocktail.

## Data Availability

The phage cocktail data presented in this study are openly available in NCBI under the following accession numbers, as described in the original papers [8,27]: ERR2184199, ERR2184200, ERR2184201, SRR3744915. The individually sequenced phage data presented in this study have been submitted to NCBI and can be found under the GenBank accession numbers OQ338181 - OQ338185, and the sequencing data can be found under the accession number SRR23151975 - SRR23151979. Code can be found at https://github.com/deng-lab/ProphageCocktail

## Acknowledgments

The authors would like to thank Dr. Hans and Prof. Peschel for kindly providing the bacterial strains used in this study. The authors also thank Sarah Pfeifer-Nigisch, Monique Preusse and Silke Bernhöft for their technical assistance. This work was funded by the German Research Foundation (D.F.G. Emmy Noether program, Project No. 273124240, SFB 1371, Project No. 395357507), Marie Sklodowska-Curie Actions Innovation Training Networks grant agreement no. 955974(VIROINF), and the European Research Council Starting grant (ERC StG 803077) awarded to L.D.

## Author Contributions

S.E.S. and W.H. isolated phages; S.E.S and X.P drafted the manuscript; X.P. performed the analyses; J.R. contributed to the analyses; M.K.M. and L.D. conceived and supervised the study and revised the manuscript. All authors reviewed and approved the manuscript.

## Notes

### Competing Interest Statement

The authors have declared no competing interest.

