## Supplemental Files for "Metagenomic analyses of single phages and phage cocktails show instances of contamination with temperate phages and bacterial DNA"

| #FILE | NAME | SEQUENCE | START | END | STRAND | GENE | COVERAGE | COVERAGE_MAP | GAPS | %COVERAGE | %IDENTITY | DATABASE | ACCESSION | PRODUCT | RESISTANCE |
| --- | --- | --- | --- | --- | --- | --- | --- | --- | --- | --- | --- | --- | --- | --- | --- |
| merged.all.fna | Staphylococcus phage HMGUsa2 | SAP2_NODE_14_length_3665_cov_4.302216 | 2044 | 3573 | - | aur | 1-1530/1530 | =============== | 0/0 | 100.00 | 99.61 | vfdb | NP_647375 | (aur) zinc metalloproteinase aureolysin [Aureolysin (VF0024)] [Staphylococcus aureus subsp. aureus MW2] | |
| merged.all.fna | Staphylococcus phage HMGUsa3 | SAP2_NODE_34_length_2263_cov_3.186141 | 131 | 436 | - | icaD | 1-306/306 | =============== | 0/0 | 100.00 | 100.00 | vfdb | NP_647404 | (icaD) intercellular adhesion protein D involved in polysaccharide intercellular adhesin(PIA) synthesis [Intercellular adhesion proteins (VF0014)] [Staphylococcus aureus subsp. aureus MW2] | |
| merged.all.fna | Staphylococcus phage HMGUsa4 | SAP2_NODE_34_length_2263_cov_3.186141 | 400 | 1638 | - | icaA | 1-1239/1239 | =============== | 0/0 | 100.00 | 99.92 | vfdb | NP_647403 | (icaA) N-acetylglucosaminyltransferase involved in polysaccharide intercellular adhesin(PIA) synthesis [Intercellular adhesion proteins (VF0014)] [Staphylococcus aureus subsp. aureus MW2] | |
| merged.all.fna | Staphylococcus phage HMGUsa5 | SAP2_NODE_34_length_2263_cov_3.186141 | 1802 | 2263 | + | icaR | 1-462/561 | =============.. | 0/0 | 82.35 | 100.00 | vfdb | NP_647402 | (icaR) ica operon transcriptional regulator IcaR [Intercellular adhesion proteins (VF0014)] [Staphylococcus aureus subsp. aureus MW2] | |
| merged.all.fna | Staphylococcus phage HMGUsa6 | SAP2_NODE_51_length_1675_cov_2.744444 | 112 | 628 | - | cap8A | 1-516/516 | ========/====== | 01-Jan | 100.00 | 99.42 | vfdb | NP_644939 | (cap8A) capsular polysaccharide synthesis enzyme [Capsule (VF0003)] [Staphylococcus aureus subsp. aureus MW2] | |
| merged.all.fna | Staphylococcus phage HMGUsa7 | SAP2_NODE_68_length_1359_cov_1.798313 | 105 | 1214 | - | cap8F | 1-1110/1110 | =============== | 0/0 | 100.00 | 99.55 | vfdb | NP_644944 | (cap8F) capsular polysaccharide synthesis enzyme Cap8F [Capsule (VF0003)] [Staphylococcus aureus subsp. aureus MW2] | |

Table S1. Virulence gene analysis

Table S2: Analysis of bacterial contamination

| idname | id | total_clean_reads | map_virus_reads_count | map_virus_rate(%) | map_bacteria_reads_count | overall_map_bacteria_rate(%) | bacteria_reference |
| --- | --- | --- | --- | --- | --- | --- | --- |
| ERR2184201 | PYO2014 | 30882170 | 15226019 | 49.3 | 233 | 0.0008 | *Staphylococcus aureus* |
| ERR2184201 | PYO2014 | 30882170 | 15226019 | 49.3 | 106 | 0.0003 | *Proteus mirabilis* |
| ERR2184199 | PYO97 | 8514396 | 4199761 | 49.33 | 1446 | 0.017 | *Escherichia coli(2)* |
| P4Ob1 | Pseudomonas phage HMGUpa1 | 11444854 | 11441751 | 99.97 | 2 | 0.0 | *Serratia marcescens* |
| ERR2184201 | PYO2014 | 30882170 | 15226019 | 49.3 | 2559 | 0.0083 | *Shigella flexneri* |
| ERR2184200 | PYO2000 | 1057238 | 165587 | 15.66 | 66 | 0.0062 | *Serratia marcescens* |
| SRR3744915 | INTESTI | 879654 | 421529 | 47.92 | 30 | 0.0034 | *Staphylococcus aureus* |
| ERR2184200 | PYO2000 | 1057238 | 165587 | 15.66 | 30 | 0.0028 | *Proteus mirabilis* |
| P4Ob1 | Pseudomonas phage HMGUpa1 | 11444854 | 11441751 | 99.97 | 0 | 0.0 | *Proteus mirabilis* |
| SM14O0610B | Serratia phage HMGUsm2 | 15312134 | 15300115 | 99.92 | 148 | 0.001 | *Shigella flexneri* |
| SAP2 | Staphylococcus phage HMGUsa2 | 12127722 | 12068558 | 99.51 | 57 | 0.0005 | *Escherichia coli(2)* |
| SAP1 | Staphylococcus phage HMGUsa1 | 9471354 | 9311859 | 98.32 | 1304 | 0.0138 | *Shigella flexneri* |
| SAP2 | Staphylococcus phage HMGUsa2 | 12127722 | 12068558 | 99.51 | 83 | 0.0007 | *Enterococcus faecium* |
| P14Ob2 | Pseudomonas phage HMGUpa2 | 16146040 | 16142069 | 99.98 | 1723 | 0.0107 | *Pseudomonas aeruginosa* |
| SM14O0610B | Serratia phage HMGUsm2 | 15312134 | 15300115 | 99.92 | 188 | 0.0012 | *Escherichia coli(2)* |
| SM14O0610B | Serratia phage HMGUsm2 | 15312134 | 15300115 | 99.92 | 117 | 0.0008 | *Pseudomonas aeruginosa* |
| SRR3744915 | INTESTI | 879654 | 421529 | 47.92 | 23 | 0.0026 | *Proteus mirabilis* |
| SAP1 | Staphylococcus phage HMGUsa1 | 9471354 | 9311859 | 98.32 | 15482 | 0.1635 | *Serratia marcescens* |
| ERR2184199 | PYO97 | 8514396 | 4199761 | 49.33 | 117 | 0.0014 | *Serratia marcescens* |
| ERR2184199 | PYO97 | 8514396 | 4199761 | 49.33 | 1284 | 0.0151 | *Shigella flexneri* |
| SAP1 | Staphylococcus phage HMGUsa1 | 9471354 | 9311859 | 98.32 | 1752 | 0.0185 | *Pseudomonas aeruginosa* |
| P14Ob2 | Pseudomonas phage HMGUpa2 | 16146040 | 16142069 | 99.98 | 17 | 0.0001 | *Enterococcus faecium* |
| SRR3744915 | INTESTI | 879654 | 421529 | 47.92 | 93 | 0.0106 | *Escherichia coli(2)* |
| SRR3744915 | INTESTI | 879654 | 421529 | 47.92 | 6 | 0.0007 | *Pseudomonas aeruginosa* |
| SM14O0610B | Serratia phage HMGUsm2 | 15312134 | 15300115 | 99.92 | 5 | 0.0 | *Enterococcus faecium* |
| SAP2 | Staphylococcus phage HMGUsa2 | 12127722 | 12068558 | 99.51 | 69 | 0.0006 | *Serratia marcescens* |
| SM14O0610B | Serratia phage HMGUsm2 | 15312134 | 15300115 | 99.92 | 4207 | 0.0275 | *Serratia marcescens* |
| ERR2184201 | PYO2014 | 30882170 | 15226019 | 49.3 | 106 | 0.0003 | *Pseudomonas aeruginosa* |
| ERR2184199 | PYO97 | 8514396 | 4199761 | 49.33 | 129 | 0.0015 | *Proteus mirabilis* |
| P4Ob1 | Pseudomonas phage HMGUpa1 | 11444854 | 11441751 | 99.97 | 4 | 0.0 | *Enterococcus faecium* |
| SAP2 | Staphylococcus phage HMGUsa2 | 12127722 | 12068558 | 99.51 | 24 | 0.0002 | *Proteus mirabilis* |
| P14Ob2 | Pseudomonas phage HMGUpa2 | 16146040 | 16142069 | 99.98 | 32 | 0.0002 | *Proteus mirabilis* |
| P14Ob2 | Pseudomonas phage HMGUpa2 | 16146040 | 16142069 | 99.98 | 73 | 0.0005 | *Serratia marcescens* |
| ERR2184199 | PYO97 | 8514396 | 4199761 | 49.33 | 53 | 0.0006 | *Pseudomonas aeruginosa* |
| ERR2184200 | PYO2000 | 1057238 | 165587 | 15.66 | 12 | 0.0011 | *Staphylococcus aureus* |
| SRR3744915 | INTESTI | 879654 | 421529 | 47.92 | 62 | 0.007 | *Shigella flexneri* |
| SAP2 | Staphylococcus phage HMGUsa2 | 12127722 | 12068558 | 99.51 | 21692 | 0.1789 | *Staphylococcus aureus* |
| SAP2 | Staphylococcus phage HMGUsa2 | 12127722 | 12068558 | 99.51 | 172 | 0.0014 | *Pseudomonas aeruginosa* |
| SRR3744915 | INTESTI | 879654 | 421529 | 47.92 | 9 | 0.001 | *Serratia marcescens* |
| SRR3744915 | INTESTI | 879654 | 421529 | 47.92 | 115 | 0.0131 | *Enterococcus faecium* |
| ERR2184199 | PYO97 | 8514396 | 4199761 | 49.33 | 40 | 0.0005 | *Staphylococcus aureus* |
| SAP1 | Staphylococcus phage HMGUsa1 | 9471354 | 9311859 | 98.32 | 2404 | 0.0254 | *Escherichia coli(2)* |
| P4Ob1 | Pseudomonas phage HMGUpa1 | 11444854 | 11441751 | 99.97 | 0 | 0.0 | *Escherichia coli(2)* |
| SAP1 | Staphylococcus phage HMGUsa1 | 9471354 | 9311859 | 98.32 | 4836 | 0.0511 | *Staphylococcus aureus* |
| P14Ob2 | Pseudomonas phage HMGUpa2 | 16146040 | 16142069 | 99.98 | 16 | 0.0001 | *Staphylococcus aureus* |
| ERR2184199 | PYO97 | 8514396 | 4199761 | 49.33 | 44 | 0.0005 | *Enterococcus faecium* |
| SAP2 | Staphylococcus phage HMGUsa2 | 12127722 | 12068558 | 99.51 | 39 | 0.0003 | *Shigella flexneri* |
| SAP1 | Staphylococcus phage HMGUsa1 | 9471354 | 9311859 | 98.32 | 189 | 0.002 | *Enterococcus faecium* |
| P4Ob1 | Pseudomonas phage HMGUpa1 | 11444854 | 11441751 | 99.97 | 0 | 0.0 | *Shigella flexneri* |
| ERR2184201 | PYO2014 | 30882170 | 15226019 | 49.3 | 3138 | 0.0102 | *Escherichia coli(2)* |
| ERR2184200 | PYO2000 | 1057238 | 165587 | 15.66 | 896 | 0.0847 | *Escherichia coli(2)* |
| ERR2184201 | PYO2014 | 30882170 | 15226019 | 49.3 | 109 | 0.0004 | *Enterococcus faecium* |
| P14Ob2 | Pseudomonas phage HMGUpa2 | 16146040 | 16142069 | 99.98 | 51 | 0.0003 | *Shigella flexneri* |
| P14Ob2 | Pseudomonas phage HMGUpa2 | 16146040 | 16142069 | 99.98 | 57 | 0.0004 | *Escherichia coli(2)* |
| P4Ob1 | Pseudomonas phage HMGUpa1 | 11444854 | 11441751 | 99.97 | 126 | 0.0011 | *Pseudomonas aeruginosa* |
| SM14O0610B | Serratia phage HMGUsm2 | 15312134 | 15300115 | 99.92 | 23 | 0.0002 | *Proteus mirabilis* |
| ERR2184200 | PYO2000 | 1057238 | 165587 | 15.66 | 7652 | 0.7238 | *Shigella flexneri* |
| P4Ob1 | Pseudomonas phage HMGUpa1 | 11444854 | 11441751 | 99.97 | 4 | 0.0 | *Staphylococcus aureus* |
| ERR2184200 | PYO2000 | 1057238 | 165587 | 15.66 | 19 | 0.0018 | *Pseudomonas aeruginosa* |
| ERR2184201 | PYO2014 | 30882170 | 15226019 | 49.3 | 273 | 0.0009 | *Serratia marcescens* |
| SAP1 | Staphylococcus phage HMGUsa1 | 9471354 | 9311859 | 98.32 | 416 | 0.0044 | *Proteus mirabilis* |
| SM14O0610B | Serratia phage HMGUsm2 | 15312134 | 15300115 | 99.92 | 2 | 0.0 | *Staphylococcus aureus* |
| ERR2184200 | PYO2000 | 1057238 | 165587 | 15.66 | 11 | 0.001 | *Enterococcus faecium* |

Table S3. Temperate contigs in cocktails. Contigs in which an integrase gene can be identified are shown with a yellow background.

| **Sample** | **reference_sequence_name** | **sequence_length** | **Relative_abundance** | **checkv_quality** | **max_score_group** | **integrase_number** | **excisionase_number** | **Replidec_label** | **bin_number** | **host_sp** | **blastn_best_hit** | **evalue** | **query cover** | **identity** |
| --- | --- | --- | --- | --- | --- | --- | --- | --- | --- | --- | --- | --- | --- | --- |
| ERR2184201 | NODE_14_length_43277_cov_21.057205 | 43277 | 0.0179 | Medium-quality | dsDNAphage | **1** | 0 | Temperate | unknown | s__Pseudomonas aeruginosa | *Pseudomonas aeruginosa* *CP082822* | 0 | 47% | 96.63 |
| ERR2184201 | NODE_19_length_39338_cov_10.558220 | 39338 | 0.0088 | Medium-quality | dsDNAphage | **1** | 0 | Temperate | ERR2184201_40 | s__Enterococcus_B faecium | Enterococcus faecium CP047326.1 | 0 | 65% | 96.68 |
| ERR2184201 | NODE_96_length_6777_cov_430.809075 | 6777 | 0.364 | Low-quality | ssDNA | **1** | 0 | Temperate | unknown | s__Escherichia coli | Escherichia coli CP002729.1 | 0 | 100% | 99.44 |
| ERR2184201 | NODE_305_length_2326_cov_2.149978 | 2326 | 0.0019 | Low-quality | ssDNA | **1** | 0 | Temperate | ERR2184201_36 | unknown | - | - | - | - |
| ERR2184199 | NODE_65_length_7075_cov_6.982621 | 7075 | 0.0135 | Low-quality | dsDNAphage | **1** | 0 | Temperate | ERR2184199_25 | s__Proteus mirabilis | Proteus mirabilis CP68152.1 | 0 | 71% | 98.76 |
| ERR2184199 | NODE_402_length_1157_cov_2.213249 | 1157 | 0.0038 | Low-quality | ssDNA | **1** | 0 | Temperate | unknown | s__Proteus mirabilis | Proteus mirabilis CP043332.1 | 0 | 99% | 99.15 |
| ERR2184200 | NODE_102_length_1807_cov_1.829040 | 1807 | 0.0194 | Low-quality | dsDNAphage | 0 | 0 | Temperate | unknown | s__Pseudomonas aeruginosa |  |  |  |  |
| ERR2184200 | NODE_129_length_1590_cov_1.945003 | 1590 | 0.0238 | Low-quality | dsDNAphage | 0 | 0 | Temperate | unknown | s__Pseudomonas aeruginosa |  |  |  |  |
| ERR2184200 | NODE_132_length_1553_cov_1.952545 | 1553 | 0.0209 | Low-quality | ssDNA | 0 | 0 | Temperate | unknown | s__Pseudomonas aeruginosa |  |  |  |  |
| ERR2184200 | NODE_139_length_1491_cov_2.640805 | 1491 | 0.0308 | Low-quality | ssDNA | 0 | 0 | Temperate | unknown | s__Pseudomonas aeruginosa |  |  |  |  |
| ERR2184201 | NODE_5_length_81730_cov_107.739082 | 81730 | 0.09 | Low-quality | dsDNAphage | 0 | 0 | Temperate | unknown | unknown |  |  |  |  |
| ERR2184201 | NODE_22_length_38163_cov_96.063288 | 38163 | 0.0804 | Low-quality | dsDNAphage | 0 | 0 | Temperate | ERR2184201_29 | unknown |  |  |  |  |
| ERR2184201 | NODE_56_length_10045_cov_4.125276 | 10045 | 0.0034 | Low-quality | dsDNAphage | 0 | 0 | Temperate | unknown | unknown |  |  |  |  |
| ERR2184201 | NODE_61_length_9837_cov_3.209694 | 9837 | 0.0028 | Low-quality | dsDNAphage | 0 | 0 | Temperate | unknown | unknown |  |  |  |  |
| ERR2184201 | NODE_102_length_6484_cov_2.469225 | 6484 | 0.0021 | Low-quality | dsDNAphage | 0 | 0 | Temperate | ERR2184201_32 | unknown |  |  |  |  |
| ERR2184201 | NODE_108_length_5835_cov_2.929916 | 5835 | 0.0025 | Low-quality | dsDNAphage | 0 | 0 | Temperate | ERR2184201_31 | unknown |  |  |  |  |
| ERR2184201 | NODE_128_length_5106_cov_2.035550 | 5106 | 0.0018 | Low-quality | dsDNAphage | 0 | 0 | Temperate | unknown | unknown |  |  |  |  |
| ERR2184201 | NODE_193_length_3556_cov_2.741105 | 3556 | 0.0023 | Low-quality | ssDNA | 0 | 0 | Temperate | ERR2184201_31 | unknown |  |  |  |  |
| ERR2184201 | NODE_195_length_3519_cov_2.770468 | 3519 | 0.0024 | Low-quality | dsDNAphage | 0 | 0 | Temperate | ERR2184201_31 | unknown |  |  |  |  |
| ERR2184201 | NODE_221_length_3180_cov_2.858812 | 3180 | 0.0024 | Low-quality | dsDNAphage | 0 | 0 | Temperate | unknown | s__Staphylococcus aureus |  |  |  |  |
| ERR2184201 | NODE_242_length_2916_cov_2.544906 | 2916 | 0.0021 | Low-quality | ssDNA | 0 | 0 | Temperate | ERR2184201_32 | unknown |  |  |  |  |
| ERR2184201 | NODE_255_length_2737_cov_1.813874 | 2737 | 0.0018 | Low-quality | dsDNAphage | 0 | 0 | Temperate | unknown | s__Vibrio splendidus_D |  |  |  |  |
| ERR2184201 | NODE_257_length_2726_cov_2.341073 | 2726 | 0.002 | Low-quality | dsDNAphage | 0 | 0 | Temperate | ERR2184201_36 | unknown |  |  |  |  |
| ERR2184201 | NODE_277_length_2571_cov_2.196602 | 2571 | 0.0017 | Low-quality | dsDNAphage | 0 | 0 | Temperate | ERR2184201_37 | unknown |  |  |  |  |
| ERR2184201 | NODE_288_length_2448_cov_2.076628 | 2448 | 0.0018 | Low-quality | dsDNAphage | 0 | 0 | Temperate | unknown | s__Staphylococcus aureus |  |  |  |  |
| ERR2184201 | NODE_339_length_2085_cov_2.251762 | 2085 | 0.0017 | Low-quality | dsDNAphage | 0 | 0 | Temperate | ERR2184201_39 | unknown |  |  |  |  |
| ERR2184201 | NODE_369_length_1886_cov_1.881365 | 1886 | 0.0015 | Low-quality | dsDNAphage | 0 | 0 | Temperate | unknown | s__Escherichia coli_C |  |  |  |  |
| ERR2184201 | NODE_381_length_1806_cov_8.439953 | 1806 | 0.01 | Low-quality | ssDNA | 0 | 0 | Temperate | unknown | unknown |  |  |  |  |
| ERR2184201 | NODE_399_length_1751_cov_1.667676 | 1751 | 0.0014 | Low-quality | dsDNAphage | 0 | 0 | Temperate | unknown | s__Escherichia coli_C |  |  |  |  |
| ERR2184201 | NODE_413_length_1699_cov_46.421875 | 1699 | 0.0441 | Low-quality | ssDNA | 0 | 0 | Temperate | unknown | unknown |  |  |  |  |
| ERR2184201 | NODE_420_length_1671_cov_2.129135 | 1671 | 0.0016 | Low-quality | dsDNAphage | 0 | 0 | Temperate | unknown | unknown |  |  |  |  |
| ERR2184201 | NODE_451_length_1570_cov_2.710401 | 1570 | 0.0021 | Low-quality | ssDNA | 0 | 0 | Temperate | unknown | s__Staphylococcus aureus |  |  |  |  |
| ERR2184201 | NODE_492_length_1468_cov_1.924763 | 1468 | 0.0017 | Low-quality | ssDNA | 0 | 0 | Temperate | unknown | s__Escherichia fergusonii |  |  |  |  |
| ERR2184201 | NODE_502_length_1455_cov_2.033186 | 1455 | 0.0017 | Low-quality | dsDNAphage | 0 | 0 | Temperate | unknown | unknown |  |  |  |  |
| ERR2184201 | NODE_525_length_1423_cov_1.338369 | 1423 | 0.0011 | Low-quality | dsDNAphage | 0 | 0 | Temperate | unknown | s__Staphylococcus aureus |  |  |  |  |
| ERR2184201 | NODE_537_length_1405_cov_1.428025 | 1405 | 0.0011 | Low-quality | dsDNAphage | 0 | 0 | Temperate | unknown | s__ |  |  |  |  |
| ERR2184201 | NODE_538_length_1404_cov_2.059770 | 1404 | 0.0015 | Low-quality | dsDNAphage | 0 | 0 | Temperate | unknown | unknown |  |  |  |  |
| ERR2184201 | NODE_565_length_1329_cov_1.713821 | 1329 | 0.0016 | Low-quality | dsDNAphage | 0 | 0 | Temperate | unknown | s__Staphylococcus aureus |  |  |  |  |
| ERR2184201 | NODE_573_length_1291_cov_1.505034 | 1291 | 0.0012 | Low-quality | ssDNA | 0 | 0 | Temperate | unknown | unknown |  |  |  |  |
| ERR2184201 | NODE_595_length_1242_cov_1.191601 | 1242 | 0.0009 | Low-quality | dsDNAphage | 0 | 0 | Temperate | unknown | s__Staphylococcus aureus |  |  |  |  |
| ERR2184201 | NODE_620_length_1197_cov_1.125683 | 1197 | 0.0009 | Low-quality | dsDNAphage | 0 | 0 | Temperate | unknown | s__Ochrobactrum intermedium | |  |  |  |
| ERR2184201 | NODE_636_length_1166_cov_1.510778 | 1166 | 0.0013 | Low-quality | dsDNAphage | 0 | 0 | Temperate | unknown | s__Staphylococcus aureus |  |  |  |  |
| ERR2184201 | NODE_650_length_1136_cov_1.783992 | 1136 | 0.0016 | Low-quality | dsDNAphage | 0 | 0 | Temperate | unknown | s__Staphylococcus aureus |  |  |  |  |
| ERR2184201 | NODE_653_length_1134_cov_1.400966 | 1134 | 0.0011 | Low-quality | dsDNAphage | 0 | 0 | Temperate | unknown | s__Staphylococcus aureus |  |  |  |  |
| ERR2184201 | NODE_658_length_1123_cov_1.357422 | 1123 | 0.0011 | Low-quality | ssDNA | 0 | 0 | Temperate | unknown | s__Psychrobacter sp001652315 | |  |  |  |
| ERR2184201 | NODE_680_length_1097_cov_7.190381 | 1097 | 0.0059 | Low-quality | dsDNAphage | 0 | 0 | Temperate | unknown | unknown |  |  |  |  |
| ERR2184201 | NODE_696_length_1074_cov_1.301538 | 1074 | 0.001 | Low-quality | dsDNAphage | 0 | 0 | Temperate | unknown | s__Staphylococcus aureus |  |  |  |  |
| ERR2184201 | NODE_710_length_1052_cov_1.541448 | 1052 | 0.0013 | Low-quality | dsDNAphage | 0 | 0 | Temperate | unknown | s__Staphylococcus aureus |  |  |  |  |
| ERR2184201 | NODE_726_length_1029_cov_1.923656 | 1029 | 0.0013 | Low-quality | dsDNAphage | 0 | 0 | Temperate | unknown | unknown |  |  |  |  |
| ERR2184201 | NODE_746_length_1010_cov_19.429199 | 1010 | 0.0179 | Low-quality | ssDNA | 0 | 0 | Temperate | unknown | unknown |  |  |  |  |
| ERR2184201 | NODE_755_length_1000_cov_1.385128 | 1000 | 0.0011 | Low-quality | dsDNAphage | 0 | 0 | Temperate | unknown | s__Staphylococcus aureus |  |  |  |  |
| SRR3744915 | NODE_4_length_87952_cov_13.069354 | 87952 | 0.2254 | Medium-quality | dsDNAphage | 0 | 0 | Temperate | SRR3744915_12 | unknown |  |  |  |  |
| SRR3744915 | NODE_14_length_39533_cov_13.474543 | 39533 | 0.2288 | Low-quality | dsDNAphage | 0 | 0 | Temperate | SRR3744915_12 | unknown |  |  |  |  |
| SRR3744915 | NODE_42_length_7788_cov_18.328721 | 7788 | 0.3048 | Low-quality | dsDNAphage | 0 | 0 | Temperate | unknown | unknown |  |  |  |  |
| SRR3744915 | NODE_61_length_4505_cov_23.222247 | 4505 | 0.4097 | Low-quality | dsDNAphage | 0 | 0 | Temperate | unknown | s__Prevotella amnii |  |  |  |  |
| SRR3744915 | NODE_85_length_3290_cov_3.054714 | 3290 | 0.0446 | Low-quality | dsDNAphage | 0 | 0 | Temperate | unknown | s__Enterococcus_B lactis |  |  |  |  |
| SRR3744915 | NODE_108_length_2540_cov_4.763783 | 2540 | 0.0747 | Low-quality | dsDNAphage | 0 | 0 | Temperate | SRR3744915_2 | s__Enterococcus_B faecium |  |  |  |  |
| SRR3744915 | NODE_199_length_1123_cov_3.288390 | 1123 | 0.0576 | Low-quality | dsDNAphage | 0 | 0 | Temperate | unknown | s__Enterococcus_B faecium |  |  |  |  |
| SRR3744915 | NODE_212_length_1047_cov_2.756048 | 1047 | 0.0453 | Low-quality | dsDNAphage | 0 | 0 | Temperate | unknown | s__Enterococcus_B lactis |  |  |  |  |
| SRR3744915 | NODE_224_length_1003_cov_24.828059 | 1003 | 0.4106 | Low-quality | ssDNA | 0 | 0 | Temperate | unknown | unknown |  |  |  |  |
| ERR2184199 | NODE_26_length_30342_cov_7.389870 | 30342 | 0.0146 | Medium-quality | dsDNAphage | 0 | 0 | Temperate | ERR2184199_25 | s__Proteus sp003144395 |  |  |  |  |
| ERR2184199 | NODE_34_length_23295_cov_4.838296 | 23295 | 0.0095 | Low-quality | dsDNAphage | 0 | 0 | Temperate | ERR2184199_19 | s__Enterococcus faecalis |  |  |  |  |
| ERR2184199 | NODE_96_length_4676_cov_2.429561 | 4676 | 0.0048 | Low-quality | dsDNAphage | 0 | 0 | Temperate | unknown | s__Pseudomonas aeruginosa |  |  |  |  |
| ERR2184199 | NODE_97_length_4553_cov_2.803913 | 4553 | 0.006 | Low-quality | dsDNAphage | 0 | 0 | Temperate | unknown | s__Enterococcus faecalis |  |  |  |  |
| ERR2184199 | NODE_116_length_3848_cov_2.427893 | 3848 | 0.005 | Low-quality | dsDNAphage | 0 | 0 | Temperate | unknown | s__Escherichia flexneri |  |  |  |  |
| ERR2184199 | NODE_137_length_3361_cov_3.679976 | 3361 | 0.007 | Low-quality | dsDNAphage | 0 | 0 | Temperate | ERR2184199_35 | s__Enterococcus_B durans |  |  |  |  |
| ERR2184199 | NODE_153_length_2956_cov_3.206825 | 2956 | 0.0061 | Low-quality | ssDNA | 0 | 0 | Temperate | unknown | s__Proteus sp001722135 |  |  |  |  |
| ERR2184199 | NODE_161_length_2761_cov_2.614930 | 2761 | 0.0047 | Low-quality | dsDNAphage | 0 | 0 | Temperate | unknown | unknown |  |  |  |  |
| ERR2184199 | NODE_202_length_2256_cov_3.890959 | 2256 | 0.0069 | Low-quality | ssDNA | 0 | 0 | Temperate | unknown | s__Proteus sp001722135 |  |  |  |  |
| ERR2184199 | NODE_210_length_2187_cov_5.233114 | 2187 | 0.0111 | Low-quality | dsDNAphage | 0 | 0 | Temperate | ERR2184199_19 | s__Proteus sp001722135 |  |  |  |  |
| ERR2184199 | NODE_228_length_2050_cov_2.133333 | 2050 | 0.0044 | Low-quality | ssDNA | 0 | 0 | Temperate | unknown | s__Escherichia coli |  |  |  |  |
| ERR2184199 | NODE_255_length_1834_cov_2.486228 | 1834 | 0.0045 | Low-quality | dsDNAphage | 0 | 0 | Temperate | unknown | s__Proteus vulgaris |  |  |  |  |
| ERR2184199 | NODE_265_length_1723_cov_1.389688 | 1723 | 0.0029 | Low-quality | dsDNAphage | 0 | 0 | Temperate | unknown | s__Proteus mirabilis |  |  |  |  |
| ERR2184199 | NODE_266_length_1715_cov_2.438554 | 1715 | 0.0049 | Low-quality | ssDNA | 0 | 0 | Temperate | unknown | s__Escherichia coli |  |  |  |  |
| ERR2184199 | NODE_270_length_1660_cov_3.146417 | 1660 | 0.0054 | Low-quality | dsDNAphage | 0 | 0 | Temperate | unknown | s__Proteus mirabilis |  |  |  |  |
| ERR2184199 | NODE_271_length_1649_cov_2.689460 | 1649 | 0.0046 | Low-quality | dsDNAphage | 0 | 0 | Temperate | unknown | s__Proteus mirabilis |  |  |  |  |
| ERR2184199 | NODE_278_length_1615_cov_1.664744 | 1615 | 0.0034 | Low-quality | ssDNA | 0 | 0 | Temperate | unknown | s__Escherichia coli |  |  |  |  |
| ERR2184199 | NODE_279_length_1613_cov_1.311297 | 1613 | 0.0022 | Low-quality | dsDNAphage | 0 | 0 | Temperate | unknown | s__Bacillus_A anthracis |  |  |  |  |
| ERR2184199 | NODE_304_length_1470_cov_1.703180 | 1470 | 0.0032 | Low-quality | dsDNAphage | 0 | 0 | Temperate | unknown | s__Pseudomonas aeruginosa |  |  |  |  |
| ERR2184199 | NODE_320_length_1399_cov_2.441964 | 1399 | 0.0047 | Low-quality | ssDNA | 0 | 0 | Temperate | unknown | s__Bacillus_A cereus |  |  |  |  |
| ERR2184199 | NODE_343_length_1330_cov_1.935686 | 1330 | 0.0035 | Low-quality | dsDNAphage | 0 | 0 | Temperate | unknown | s__Escherichia coli_D |  |  |  |  |
| ERR2184199 | NODE_346_length_1326_cov_1.571204 | 1326 | 0.003 | Low-quality | dsDNAphage | 0 | 0 | Temperate | unknown | s__Bacillus_A anthracis |  |  |  |  |
| ERR2184199 | NODE_353_length_1308_cov_2.289705 | 1308 | 0.0044 | Low-quality | dsDNAphage | 0 | 0 | Temperate | unknown | s__Bacillus_A proteolyticus |  |  |  |  |
| ERR2184199 | NODE_367_length_1258_cov_2.975062 | 1258 | 0.0058 | Low-quality | dsDNAphage | 0 | 0 | Temperate | unknown | s__Escherichia coli_D |  |  |  |  |
| ERR2184199 | NODE_381_length_1219_cov_2.091065 | 1219 | 0.0039 | Low-quality | ssDNA | 0 | 0 | Temperate | unknown | s__Proteus sp003144395 |  |  |  |  |
| ERR2184199 | NODE_399_length_1165_cov_1.845045 | 1165 | 0.0034 | Low-quality | dsDNAphage | 0 | 0 | Temperate | unknown | unknown |  |  |  |  |
| ERR2184199 | NODE_415_length_1123_cov_2.023408 | 1123 | 0.0029 | Low-quality | ssDNA | 0 | 0 | Temperate | unknown | s__Pseudomonas aeruginosa |  |  |  |  |
| ERR2184199 | NODE_441_length_1054_cov_2.570571 | 1054 | 0.0048 | Low-quality | ssDNA | 0 | 0 | Temperate | unknown | s__Proteus mirabilis |  |  |  |  |
| ERR2184199 | NODE_469_length_1001_cov_1.381607 | 1001 | 0.0025 | Low-quality | dsDNAphage | 0 | 0 | Temperate | unknown | s__Enterococcus_B faecium |  |  |  |  |

Table S4. Bacterial strains used in this study.

| **Label** | **Strain ID** | **Species** | **Source** |
| --- | --- | --- | --- |
| PAO1 | PAO1 | *P. aeuruginosa* | Dr. Michael Rothballer (Helmholtz Zentrum München, Germany) |
| 1E | 073521-1E, clinical isolate | *P. aeuruginosa* |  |
| 50071 | DSMZ 50071 | *P. aeuruginosa* |  |
| P1 | NMI 3816/05 (PL3816) | *P. aeuruginosa* | Paweł Urbanowicz (Department of Molecular Microbiology, National Medicines Institute, Poland) |
| P2 | NMI 3220/05 (PL3220) | *P. aeuruginosa* |  |
| P3 | NMI 2427/06 (PL2427) | *P. aeuruginosa* |  |
| P4 | NMI 2409/06 (PL2409) | *P. aeuruginosa* |  |
| P5 | NMI 2312/06 (PL2312) | *P. aeuruginosa* |  |
| P6 | NMI 1186/06 (PL1186) | *P. aeuruginosa* |  |
| P7 | NMI 1025/06 (PL1025) | *P. aeuruginosa* |  |
| P8 | NMI 1010/06 (PL1010) | *P. aeuruginosa* |  |
| P9 | NMI 4992/09 | *P. aeuruginosa* |  |
| P10 | NMI 5282/13 | *P. aeuruginosa* |  |
| P11 | NMI 2260/13 | *P. aeuruginosa* |  |
| P12 | NMI 2142/13 | *P. aeuruginosa* |  |
| P13 | NMI 1334/14 | *P. aeuruginosa* |  |
| P14 | NMI 2256/14 | *P. aeuruginosa* |  |
| SA1 | N315 | *S. aureus* | Prof. Andreas Peschel (University of Tübingen, Germany) |
| SA2 | RN1 | *S. aureus* |  |
| SA3 | SA27660 | *S. aureus* |  |
| SA4 | USA300 | *S. aureus* |  |
| SA5 | SA20231 | *S. aureus* |  |
| SA6 | JS5395 | *S. aureus* |  |
| SA7 | 2220231 | *S. aureus* |  |
| SA8 | MW2 | *S. aureus* |  |
| SA9 | DSM 13661 | *S. aureus* | DSMZ |
| SA10 | DSM 779 | *S. aureus* |  |
| SA11 | Newman | *S. aureus* | Prof. Andreas Peschel (University of Tübingen, Germany) |
| SA12 | RN4220 | *S. aureus* |  |
| SA13 | PS187 | *S. aureus* |  |
| SA14 | COL | *S. aureus* |  |
| SA15 | SA113 | *S. aureus* |  |
| SA16 | Mu50 | *S. aureus* |  |
| SA17 | SH1000 | *S. aureus* |  |
| SM1 | NRZ-49082 | *S. marcescens* | Dr. Jörg B. Hans (RUHR University, Germany) |
| SM2 | NRZ-51623 | *S. marcescens* |  |
| SM3 | NRZ-38829 | *S. marcescens* |  |
| SM4 | NRZ-51155 | *S. marcescens* |  |
| SM5 | NRZ-47931 | *S. marcescens* |  |
| SM6 | NRZ-58729 | *S. marcescens* |  |
| SM7 | NRZ-20916 | *S. marcescens* |  |
| SM8 | NRZ-26344 | *S. marcescens* |  |
| SM9 | NRZ-53046 | *S. marcescens* |  |
| SM10 | NRZ-28540 | *S. marcescens* |  |
| SM11 | NRZ-28542 | *S. marcescens* |  |
| SM12 | NRZ-26667 | *S. marcescens* |  |
| SM13 | NRZ-23661 | *S. marcescens* |  |
| SM14 | NRZ-47541 | *S. marcescens* |  |
| SM15 | NRZ-56321 | *S. marcescens* |  |
| SM16 | NRZ-60428 | *S. marcescens* |  |
| SM17 | NRZ-49541 | *S. marcescens* |  |
| SM18 | NRZ-59007 | *S. marcescens* |  |
| SMC1 | sema092304 | *S. marcescens* | Clinical Isolates |
| SMC2 | sema092561 | *S. marcescens* |  |
| SMC3 | sema092193 | *S. marcescens* |  |

Table S5: mock phage cocktail

| sample_name | contig_length | provirus | gene_count | viral_genes | host_genes | checkv_quality | miuvig_quality | completeness_method | contamination | kmer_freq | warnings | integrase_number | excisionase_number | final_label | match_gene_number |
| --- | --- | --- | --- | --- | --- | --- | --- | --- | --- | --- | --- | --- | --- | --- | --- |
| **NODE_1_length_142648_cov_78.839705** | **142648** | **No** | **222** | **200** | **0** | **Complete** | **High-quality** | **DTR (high-confidence)** | **0.0** | **1.0** |  | **0.0** | **0.0** | **Virulent** | **220.0** |
| **NODE_2_length_45389_cov_81.608815** | **45389** | **No** | **68** | **46** | **0** | **Complete** | **High-quality** | **DTR (medium-confidence)** | **0.0** | **1.0** |  | **0.0** | **0.0** | **Virulent** | **66.0** |
| **NODE_3_length_43330_cov_83.078914** | **43330** | **No** | **54** | **37** | **0** | **Medium-quality** | **Genome-fragment** | **HMM-based (lower-bound)** | **0.0** | **1.0** | **low-confidence DTR** | **0.0** | **0.0** | **Virulent** | **54.0** |
| **NODE_4_length_39100_cov_121.022359** | **39100** | **No** | **48** | **45** | **0** | **Complete** | **High-quality** | **DTR (high-confidence)** | **0.0** | **1.0** |  | **0.0** | **0.0** | **Virulent** | **47.0** |
| **NODE_5_length_35297_cov_77.938709** | **35297** | **No** | **47** | **30** | **1** | **Medium-quality** | **Genome-fragment** | **HMM-based (lower-bound)** | **0.0** | **1.0** |  | **1.0** | **0.0** | **Temperate** | **43.0** |
| **NODE_6_length_32498_cov_78.823968** | **32498** | **No** | **40** | **33** | **1** | **Medium-quality** | **Genome-fragment** | **HMM-based (lower-bound)** | **0.0** | **1.0** |  | **1.0** | **0.0** | **Temperate** | **39.0** |
| **NODE_7_length_30757_cov_78.940330** | **30757** | **No** | **63** | **53** | **0** | **Medium-quality** | **Genome-fragment** | **HMM-based (lower-bound)** | **0.0** | **1.0** |  | **1.0** | **0.0** | **Temperate** | **63.0** |
| **NODE_8_length_27421_cov_6.662209** | **27421** | **No** | **25** | **0** | **20** | **Not-determined** | **Genome-fragment** |  | **0.0** | **1.0** | **no viral genes detected** | **1.0** | **0.0** | **Temperate** | **17.0** |
| **NODE_9_length_13934_cov_81.122415** | **13934** | **No** | **11** | **10** | **0** | **Low-quality** | **Genome-fragment** | **HMM-based (lower-bound)** | **0.0** | **1.0** |  | **0.0** | **0.0** | **Temperate** | **11.0** |
| **NODE_10_length_11199_cov_93.749551** | **11199** | **No** | **12** | **2** | **0** | **Low-quality** | **Genome-fragment** | **HMM-based (lower-bound)** | **0.0** | **1.0** |  | **1.0** | **0.0** | **Temperate** | **12.0** |
| **NODE_11_length_11142_cov_5.611617** | **11142** | **No** | **18** | **17** | **0** | **Low-quality** | **Genome-fragment** | **HMM-based (lower-bound)** | **0.0** | **1.0** |  | **0.0** | **0.0** | **Temperate** | **18.0** |
| **NODE_12_length_10819_cov_91.740338** | **10819** | **No** | **12** | **2** | **0** | **Low-quality** | **Genome-fragment** | **HMM-based (lower-bound)** | **0.0** | **1.0** |  | **1.0** | **0.0** | **Temperate** | **10.0** |
| NODE_13_length_7486_cov_4.570314 | 7486 | No | 11 | 0 | 5 | Not-determined | Genome-fragment |  | 0.0 | 1.0 | no viral genes detected | 1.0 | 0.0 | Temperate | 7.0 |
| NODE_14_length_5529_cov_5.540190 | 5529 | No | 7 | 7 | 0 | Low-quality | Genome-fragment | HMM-based (lower-bound) | 0.0 | 1.0 |  | 0.0 | 0.0 | Temperate | 7.0 |
| NODE_15_length_5150_cov_8.792738 | 5150 | No | 7 | 0 | 5 | Not-determined | Genome-fragment |  | 0.0 | 1.0 | no viral genes detected | 0.0 | 0.0 | Temperate | 4.0 |
| NODE_16_length_3831_cov_12.772246 | 3831 | No | 4 | 0 | 2 | Not-determined | Genome-fragment |  | 0.0 | 1.0 | no viral genes detected | 0.0 | 0.0 | Temperate | 1.0 |
| NODE_17_length_3810_cov_5.362716 | 3810 | No | 4 | 0 | 2 | Not-determined | Genome-fragment |  | 0.0 | 1.0 | no viral genes detected | 0.0 | 0.0 | Temperate | 2.0 |
| NODE_18_length_3678_cov_4.551753 | 3678 | No | 5 | 0 | 4 | Not-determined | Genome-fragment |  | 0.0 | 1.0 | no viral genes detected | 0.0 | 0.0 | Temperate | 1.0 |
| NODE_19_length_3449_cov_4.602534 | 3449 | No | 2 | 0 | 0 | Not-determined | Genome-fragment |  | 0.0 | 1.0 | no viral genes detected | 0.0 | 0.0 | Temperate | 2.0 |
| NODE_20_length_3446_cov_9.484223 | 3446 | No | 5 | 0 | 1 | Not-determined | Genome-fragment |  | 0.0 | 1.0 | no viral genes detected | 1.0 | 0.0 | Temperate | 4.0 |
| NODE_21_length_3361_cov_6.166062 | 3361 | No | 2 | 2 | 0 | Low-quality | Genome-fragment | HMM-based (lower-bound) | 0.0 | 1.0 |  | 0.0 | 0.0 | Temperate | 2.0 |
| NODE_22_length_3121_cov_27.022179 | 3121 | No | 1 | 0 | 0 | Not-determined | Genome-fragment |  | 0.0 | 1.0 | no viral genes detected | 0.0 | 0.0 | Temperate | 1.0 |
| NODE_23_length_3037_cov_4.022133 | 3037 | No | 5 | 0 | 2 | Not-determined | Genome-fragment |  | 0.0 | 1.0 | no viral genes detected | 0.0 | 0.0 | Temperate | 3.0 |
| NODE_24_length_2958_cov_4.944196 | 2958 | No | 4 | 0 | 4 | Not-determined | Genome-fragment |  | 0.0 | 1.0 | no viral genes detected | 0.0 | 0.0 | Temperate | 4.0 |
| NODE_25_length_2875_cov_4.252128 | 2875 | No | 2 | 2 | 0 | Low-quality | Genome-fragment | HMM-based (lower-bound) | 0.0 | 1.0 |  | 0.0 | 0.0 | Temperate | 2.0 |
| NODE_26_length_2647_cov_4.152778 | 2647 | No | 2 | 0 | 0 | Not-determined | Genome-fragment |  | 0.0 | 1.0 | no viral genes detected | 1.0 | 0.0 | Temperate | 2.0 |
| NODE_27_length_2578_cov_4.461356 | 2578 | No | 2 | 0 | 2 | Not-determined | Genome-fragment |  | 0.0 | 1.0 | no viral genes detected | 0.0 | 0.0 | Temperate | 2.0 |
| NODE_28_length_2516_cov_4.037790 | 2516 | No | 2 | 0 | 1 | Not-determined | Genome-fragment |  | 0.0 | 1.0 | no viral genes detected | 0.0 | 0.0 | Temperate | 2.0 |
| NODE_29_length_2503_cov_79.936683 | 2503 | No | 4 | 0 | 0 | Not-determined | Genome-fragment |  | 0.0 | 01-Jan | no viral genes detected; low-confidence DTR | 0.0 | 0.0 | Temperate | 1.0 |
| NODE_30_length_2418_cov_3.315277 | 2418 | No | 3 | 0 | 2 | Not-determined | Genome-fragment |  | 0.0 | 1.0 | no viral genes detected | 0.0 | 0.0 | Temperate | 1.0 |
| NODE_31_length_2321_cov_4.107237 | 2321 | No | 1 | 0 | 1 | Not-determined | Genome-fragment |  | 0.0 | 1.0 | no viral genes detected | 0.0 | 0.0 | Temperate | 1.0 |
| NODE_32_length_2313_cov_3.845438 | 2313 | No | 2 | 0 | 1 | Not-determined | Genome-fragment |  | 0.0 | 1.0 | no viral genes detected | 0.0 | 0.0 | Temperate | 2.0 |
| NODE_33_length_2275_cov_5.660811 | 2275 | No | 4 | 2 | 0 | Low-quality | Genome-fragment | HMM-based (lower-bound) | 0.0 | 1.0 |  | 0.0 | 0.0 | Temperate | 4.0 |
| NODE_34_length_2275_cov_4.045045 | 2275 | No | 4 | 0 | 0 | Not-determined | Genome-fragment |  | 0.0 | 1.0 | no viral genes detected | 0.0 | 0.0 | Temperate | 1.0 |
| NODE_35_length_2274_cov_4.441640 | 2274 | No | 3 | 3 | 0 | Low-quality | Genome-fragment | HMM-based (lower-bound) | 0.0 | 1.0 |  | 0.0 | 0.0 | Temperate | 3.0 |
| NODE_36_length_2162_cov_3.987660 | 2162 | No | 2 | 0 | 2 | Not-determined | Genome-fragment |  | 0.0 | 1.0 | no viral genes detected | 0.0 | 0.0 | Temperate | 1.0 |
| NODE_37_length_2079_cov_3.343874 | 2079 | No | 3 | 0 | 2 | Not-determined | Genome-fragment |  | 0.0 | 1.0 | no viral genes detected | 0.0 | 0.0 | Temperate | 2.0 |
| NODE_38_length_2041_cov_3.987915 | 2041 | No | 4 | 0 | 0 | Not-determined | Genome-fragment |  | 0.0 | 1.0 | no viral genes detected | 0.0 | 0.0 | Temperate | 1.0 |
| NODE_39_length_2038_cov_14.267272 | 2038 | No | 2 | 0 | 1 | Not-determined | Genome-fragment |  | 0.0 | 1.0 | no viral genes detected | 0.0 | 0.0 | Temperate | 1.0 |
| NODE_40_length_1987_cov_14.600414 | 1987 | No | 1 | 0 | 1 | Not-determined | Genome-fragment |  | 0.0 | 01-Feb | no viral genes detected; low-confidence DTR | 0.0 | 0.0 | Temperate | 1.0 |
| NODE_41_length_1877_cov_4.130626 | 1877 | No | 3 | 0 | 0 | Not-determined | Genome-fragment |  | 0.0 | 1.0 | no viral genes detected | 0.0 | 0.0 | Temperate | 3.0 |
| NODE_42_length_1747_cov_2.710993 | 1747 | No | 2 | 0 | 0 | Not-determined | Genome-fragment |  | 0.0 | 1.0 | no viral genes detected | 0.0 | 0.0 | Temperate | 2.0 |
| NODE_43_length_1729_cov_3.918757 | 1729 | No | 1 | 0 | 1 | Not-determined | Genome-fragment |  | 0.0 | 1.0 | no viral genes detected | 0.0 | 0.0 | Temperate | 1.0 |
| NODE_44_length_1719_cov_6.359375 | 1719 | No | 1 | 0 | 1 | Not-determined | Genome-fragment |  | 0.0 | 1.0 | no viral genes detected | 0.0 | 0.0 | Temperate | 1.0 |
| NODE_45_length_1670_cov_4.004954 | 1670 | No | 2 | 0 | 2 | Not-determined | Genome-fragment |  | 0.0 | 1.0 | no viral genes detected | 0.0 | 0.0 | Temperate | 2.0 |
| NODE_46_length_1641_cov_4.309584 | 1641 | No | 4 | 3 | 0 | Low-quality | Genome-fragment | HMM-based (lower-bound) | 0.0 | 1.0 |  | 0.0 | 0.0 | Temperate | 4.0 |
| NODE_47_length_1632_cov_3.247939 | 1632 | No | 2 | 0 | 1 | Not-determined | Genome-fragment |  | 0.0 | 1.0 | no viral genes detected | 0.0 | 0.0 | Temperate | 1.0 |
| NODE_48_length_1623_cov_4.832270 | 1623 | No | 1 | 1 | 0 | Low-quality | Genome-fragment | HMM-based (lower-bound) | 0.0 | 1.0 |  | 0.0 | 0.0 | Temperate | 1.0 |
| NODE_49_length_1592_cov_3.977228 | 1592 | No | 3 | 0 | 1 | Not-determined | Genome-fragment |  | 0.0 | 1.0 | no viral genes detected | 0.0 | 0.0 | Temperate | 1.0 |
| NODE_50_length_1583_cov_4.517016 | 1583 | No | 1 | 0 | 1 | Not-determined | Genome-fragment |  | 0.0 | 1.0 | no viral genes detected | 0.0 | 0.0 | Temperate | 1.0 |
| NODE_51_length_1522_cov_2.771643 | 1522 | No | 3 | 0 | 1 | Not-determined | Genome-fragment |  | 0.0 | 1.0 | no viral genes detected | 0.0 | 0.0 | Virulent | 0.0 |
| NODE_52_length_1506_cov_2.912474 | 1506 | No | 2 | 0 | 2 | Not-determined | Genome-fragment |  | 0.0 | 1.0 | no viral genes detected | 0.0 | 0.0 | Temperate | 2.0 |
| NODE_53_length_1460_cov_2.993594 | 1460 | No | 2 | 0 | 2 | Not-determined | Genome-fragment |  | 0.0 | 1.0 | no viral genes detected | 0.0 | 0.0 | Temperate | 2.0 |
| NODE_54_length_1435_cov_3.710145 | 1435 | No | 2 | 0 | 1 | Not-determined | Genome-fragment |  | 0.0 | 1.0 | no viral genes detected | 0.0 | 0.0 | Temperate | 1.0 |
| NODE_55_length_1422_cov_2.809071 | 1422 | No | 1 | 1 | 0 | Low-quality | Genome-fragment | HMM-based (lower-bound) | 0.0 | 1.0 |  | 0.0 | 0.0 | Virulent | 1.0 |
| NODE_56_length_1416_cov_3.291697 | 1416 | No | 2 | 0 | 1 | Not-determined | Genome-fragment |  | 0.0 | 1.0 | no viral genes detected | 0.0 | 0.0 | Temperate | 1.0 |
| NODE_57_length_1403_cov_3.145401 | 1403 | No | 2 | 0 | 1 | Not-determined | Genome-fragment |  | 0.0 | 1.0 | no viral genes detected | 0.0 | 0.0 | Temperate | 1.0 |
| NODE_58_length_1394_cov_5.082151 | 1394 | No | 1 | 1 | 0 | Low-quality | Genome-fragment | HMM-based (lower-bound) | 0.0 | 1.0 |  | 0.0 | 0.0 | Temperate | 1.0 |
| NODE_59_length_1375_cov_3.831818 | 1375 | No | 3 | 3 | 0 | Low-quality | Genome-fragment | HMM-based (lower-bound) | 0.0 | 1.0 |  | 0.0 | 0.0 | Temperate | 3.0 |
| NODE_60_length_1370_cov_3.224335 | 1370 | No | 1 | 0 | 0 | Not-determined | Genome-fragment |  | 0.0 | 1.0 | no viral genes detected | 0.0 | 0.0 | Temperate | 1.0 |
| NODE_61_length_1335_cov_3.625000 | 1335 | No | 3 | 0 | 0 | Not-determined | Genome-fragment |  | 0.0 | 1.0 | no viral genes detected | 0.0 | 0.0 | Temperate | 3.0 |
| NODE_62_length_1326_cov_3.713611 | 1326 | No | 3 | 0 | 1 | Not-determined | Genome-fragment |  | 0.0 | 1.0 | no viral genes detected | 0.0 | 0.0 | Temperate | 1.0 |
| NODE_63_length_1321_cov_3.464455 | 1321 | No | 2 | 0 | 1 | Not-determined | Genome-fragment |  | 0.0 | 1.0 | no viral genes detected | 0.0 | 0.0 | Temperate | 1.0 |
| NODE_64_length_1296_cov_3.828364 | 1296 | No | 2 | 1 | 0 | Low-quality | Genome-fragment | HMM-based (lower-bound) | 0.0 | 1.0 |  | 0.0 | 0.0 | Temperate | 2.0 |
| NODE_65_length_1292_cov_3.850445 | 1292 | No | 1 | 0 | 1 | Not-determined | Genome-fragment |  | 0.0 | 1.0 | no viral genes detected | 0.0 | 0.0 | Temperate | 1.0 |
| NODE_66_length_1281_cov_2.712072 | 1281 | No | 1 | 0 | 1 | Not-determined | Genome-fragment |  | 0.0 | 1.0 | no viral genes detected | 0.0 | 0.0 | Temperate | 1.0 |
| NODE_67_length_1256_cov_2.198168 | 1256 | No | 3 | 0 | 2 | Not-determined | Genome-fragment |  | 0.0 | 1.0 | no viral genes detected | 0.0 | 0.0 | Temperate | 3.0 |
| NODE_68_length_1252_cov_2.998329 | 1252 | No | 2 | 0 | 0 | Not-determined | Genome-fragment |  | 0.0 | 1.0 | no viral genes detected | 0.0 | 0.0 | Temperate | 2.0 |
| NODE_69_length_1246_cov_3.340890 | 1246 | No | 2 | 0 | 1 | Not-determined | Genome-fragment |  | 0.0 | 1.0 | no viral genes detected | 0.0 | 0.0 | Temperate | 1.0 |
| NODE_70_length_1238_cov_3.479290 | 1238 | No | 2 | 0 | 1 | Not-determined | Genome-fragment |  | 0.0 | 1.0 | no viral genes detected | 0.0 | 0.0 | Temperate | 1.0 |
| NODE_71_length_1233_cov_4.537351 | 1233 | No | 2 | 0 | 0 | Not-determined | Genome-fragment |  | 0.0 | 1.0 | no viral genes detected | 0.0 | 0.0 | Virulent | 0.0 |
| NODE_72_length_1228_cov_3.955669 | 1228 | No | 2 | 0 | 0 | Not-determined | Genome-fragment |  | 0.0 | 1.0 | no viral genes detected | 0.0 | 0.0 | Temperate | 1.0 |
| NODE_73_length_1205_cov_3.863478 | 1205 | No | 2 | 0 | 1 | Not-determined | Genome-fragment |  | 0.0 | 1.0 | no viral genes detected | 0.0 | 0.0 | Virulent | 0.0 |
| NODE_74_length_1187_cov_7.581272 | 1187 | No | 2 | 0 | 0 | Not-determined | Genome-fragment |  | 0.0 | 1.0 | no viral genes detected | 0.0 | 0.0 | Temperate | 2.0 |
| NODE_75_length_1174_cov_2.716711 | 1174 | No | 1 | 0 | 1 | Not-determined | Genome-fragment |  | 0.0 | 1.0 | no viral genes detected | 0.0 | 0.0 | Temperate | 1.0 |
| NODE_76_length_1163_cov_3.249097 | 1163 | No | 2 | 0 | 0 | Not-determined | Genome-fragment |  | 0.0 | 1.0 | no viral genes detected | 1.0 | 0.0 | Temperate | 2.0 |
| NODE_77_length_1158_cov_2.579329 | 1158 | No | 3 | 0 | 1 | Not-determined | Genome-fragment |  | 0.0 | 1.0 | no viral genes detected | 0.0 | 0.0 | Temperate | 1.0 |
| NODE_78_length_1150_cov_3.374429 | 1150 | No | 2 | 0 | 2 | Not-determined | Genome-fragment |  | 0.0 | 1.0 | no viral genes detected | 0.0 | 0.0 | Temperate | 1.0 |
| NODE_79_length_1149_cov_3.733090 | 1149 | No | 2 | 0 | 1 | Not-determined | Genome-fragment |  | 0.0 | 1.0 | no viral genes detected | 0.0 | 0.0 | Temperate | 1.0 |
| NODE_80_length_1142_cov_3.311868 | 1142 | No | 2 | 0 | 0 | Not-determined | Genome-fragment |  | 0.0 | 1.0 | no viral genes detected | 0.0 | 0.0 | Temperate | 1.0 |
| NODE_81_length_1132_cov_4.187558 | 1132 | No | 3 | 2 | 0 | Low-quality | Genome-fragment | HMM-based (lower-bound) | 0.0 | 1.0 |  | 0.0 | 0.0 | Temperate | 3.0 |
| NODE_82_length_1132_cov_3.780873 | 1132 | No | 2 | 0 | 2 | Not-determined | Genome-fragment |  | 0.0 | 1.0 | no viral genes detected | 0.0 | 0.0 | Temperate | 1.0 |
| NODE_83_length_1126_cov_4.324930 | 1126 | No | 3 | 0 | 0 | Not-determined | Genome-fragment |  | 0.0 | 1.0 | no viral genes detected | 0.0 | 0.0 | Virulent | 0.0 |
| NODE_84_length_1111_cov_3.668561 | 1111 | No | 3 | 0 | 0 | Not-determined | Genome-fragment |  | 0.0 | 1.0 | no viral genes detected | 0.0 | 0.0 | Temperate | 2.0 |
| NODE_85_length_1105_cov_3.710476 | 1105 | No | 1 | 1 | 0 | Low-quality | Genome-fragment | HMM-based (lower-bound) | 0.0 | 1.0 |  | 0.0 | 0.0 | Temperate | 1.0 |
| NODE_86_length_1099_cov_19.016284 | 1099 | No | 0 | 0 | 0 | Not-determined | Genome-fragment |  | 0.0 | 1.0 | no viral genes detected |  |  |  |  |
| NODE_87_length_1097_cov_3.432821 | 1097 | No | 1 | 0 | 1 | Not-determined | Genome-fragment |  | 0.0 | 1.0 | no viral genes detected | 0.0 | 0.0 | Temperate | 1.0 |
| NODE_88_length_1085_cov_2.593204 | 1085 | No | 2 | 0 | 0 | Not-determined | Genome-fragment |  | 0.0 | 01-Jun | no viral genes detected | 0.0 | 0.0 | Temperate | 1.0 |
| NODE_89_length_1084_cov_3.131195 | 1084 | No | 2 | 0 | 0 | Not-determined | Genome-fragment |  | 0.0 | 1.0 | no viral genes detected | 0.0 | 0.0 | Temperate | 2.0 |
| NODE_90_length_1081_cov_3.980507 | 1081 | No | 0 | 0 | 0 | Not-determined | Genome-fragment |  | 0.0 | 1.0 | no viral genes detected |  |  |  |  |
| NODE_91_length_1078_cov_3.132942 | 1078 | No | 1 | 0 | 1 | Not-determined | Genome-fragment |  | 0.0 | 1.0 | no viral genes detected | 0.0 | 0.0 | Temperate | 1.0 |
| NODE_92_length_1049_cov_3.798793 | 1049 | No | 1 | 0 | 0 | Not-determined | Genome-fragment |  | 0.0 | 1.0 | no viral genes detected | 0.0 | 0.0 | Temperate | 1.0 |
| NODE_93_length_1046_cov_4.373360 | 1046 | No | 1 | 0 | 1 | Not-determined | Genome-fragment |  | 0.0 | 1.0 | no viral genes detected | 0.0 | 0.0 | Virulent | 0.0 |
| NODE_94_length_1024_cov_3.057792 | 1024 | No | 1 | 0 | 0 | Not-determined | Genome-fragment |  | 0.0 | 1.0 | no viral genes detected | 0.0 | 0.0 | Temperate | 1.0 |
| NODE_95_length_1011_cov_2.259414 | 1011 | No | 2 | 0 | 0 | Not-determined | Genome-fragment |  | 0.0 | 1.0 | no viral genes detected | 0.0 | 0.0 | Temperate | 2.0 |
| NODE_96_length_1005_cov_3.368421 | 1005 | No | 3 | 0 | 2 | Not-determined | Genome-fragment |  | 0.0 | 1.0 | no viral genes detected | 0.0 | 0.0 | Virulent | 0.0 |
| NODE_97_length_1004_cov_2.267650 | 1004 | No | 2 | 2 | 0 | Low-quality | Genome-fragment | HMM-based (lower-bound) | 0.0 | 1.0 |  | 0.0 | 0.0 | Virulent | 2.0 |

Table S6: annotations of the temperate contigs found in the commercial cocktails

| Name | Sequence Name | Start | End | Length | Direction |
| --- | --- | --- | --- | --- | --- |
| CDS | ERR2184201_14 | 42936 | 43148 | 213 | forward |
| unknown function CDS | ERR2184201_14 | 42541 | 42903 | 363 | forward |
| unknown function CDS | ERR2184201_14 | 42312 | 42524 | 213 | forward |
| unknown function CDS | ERR2184201_14 | 41796 | 42002 | 207 | forward |
| CDS | ERR2184201_14 | 41529 | 41693 | 165 | forward |
| CDS | ERR2184201_14 | 41313 | 41516 | 204 | reverse |
| unknown function CDS | ERR2184201_14 | 40583 | 40849 | 267 | forward |
| transcriptional repressor | ERR2184201_14 | 39704 | 40573 | 870 | forward |
| CII-like regulator | ERR2184201_14 | 39280 | 39591 | 312 | reverse |
| CDS | ERR2184201_19 | 39112 | 39336 | 225 | forward |
| anti-repressor Ant | ERR2184201_14 | 38452 | 39204 | 753 | reverse |
| endolysin | ERR2184201_19 | 37933 | 38952 | 1020 | forward |
| holin | ERR2184201_19 | 37725 | 37922 | 198 | forward |
| replication initiation protein | ERR2184201_14 | 37646 | 38455 | 810 | reverse |
| holin | ERR2184201_19 | 37446 | 37709 | 264 | forward |
| unknown function CDS | ERR2184201_19 | 37294 | 37431 | 138 | forward |
| DnaC-like helicase loader | ERR2184201_14 | 36865 | 37656 | 792 | reverse |
| unknown function CDS | ERR2184201_19 | 36846 | 37292 | 447 | forward |
| unknown function CDS | ERR2184201_14 | 36180 | 36662 | 483 | reverse |
| DksA-like zinc-finger protein | ERR2184201_14 | 35978 | 36178 | 201 | reverse |
| unknown function CDS | ERR2184201_14 | 35763 | 35978 | 216 | reverse |
| NinB/ Orf homologous recombination mediator | ERR2184201_14 | 35113 | 35514 | 402 | reverse |
| CDS | ERR2184201_14 | 34862 | 35116 | 255 | reverse |
| unknown function CDS | ERR2184201_14 | 34635 | 34865 | 231 | reverse |
| unknown function CDS | ERR2184201_19 | 34476 | 36827 | 2352 | forward |
| NinG/ Rap DNA junction specific endonuclease | ERR2184201_14 | 33994 | 34638 | 645 | reverse |
| minor head protein | ERR2184201_19 | 33702 | 34460 | 759 | forward |
| anti-termination protein Q-like | ERR2184201_14 | 33623 | 33997 | 375 | reverse |
| holin | ERR2184201_14 | 33162 | 33380 | 219 | reverse |
| endolysin | ERR2184201_14 | 32713 | 33165 | 453 | reverse |
| Rz-like spanin | ERR2184201_14 | 32273 | 32716 | 444 | reverse |
| unknown function CDS | ERR2184201_14 | 32046 | 32276 | 231 | reverse |
| unknown function CDS | ERR2184201_19 | 31905 | 33689 | 1785 | forward |
| terminase small subunit | ERR2184201_14 | 31596 | 32042 | 447 | reverse |
| unknown function CDS | ERR2184201_14 | 30146 | 31615 | 1470 | reverse |
| unknown function CDS | ERR2184201_14 | 29880 | 30134 | 255 | reverse |
| single strand DNA binding protein | ERR2184199_26 | 29875 | 30342 | 468 | forward |
| exonuclease | ERR2184199_26 | 29160 | 29885 | 726 | forward |
| minor head protein | ERR2184201_19 | 29099 | 31930 | 2832 | forward |
| HNH endonuclease | ERR2184199_26 | 28571 | 29191 | 621 | forward |
| minor tail protein | ERR2184201_19 | 28314 | 29102 | 789 | forward |
| head-tail adaptor | ERR2184201_14 | 28190 | 29878 | 1689 | reverse |
| unknown function CDS | ERR2184201_14 | 27878 | 28189 | 312 | reverse |
| RecT-like ssDNA annealing protein | ERR2184199_26 | 27647 | 28531 | 885 | forward |
| CDS | ERR2184199_26 | 27447 | 27650 | 204 | forward |
| protease | ERR2184201_14 | 27261 | 27881 | 621 | reverse |
| CDS | ERR2184199_26 | 27149 | 27469 | 321 | forward |
| CDS | ERR2184199_26 | 26986 | 27141 | 156 | forward |
| unknown function CDS | ERR2184199_26 | 26580 | 26927 | 348 | forward |
| major head protein | ERR2184201_14 | 26262 | 27254 | 993 | reverse |
| unknown function CDS | ERR2184199_26 | 25960 | 26550 | 591 | forward |
| unknown function CDS | ERR2184201_14 | 25786 | 26232 | 447 | reverse |
| CDS | ERR2184199_26 | 25639 | 25950 | 312 | forward |
| unknown function CDS | ERR2184201_14 | 25533 | 25775 | 243 | reverse |
| tail length tape measure protein | ERR2184201_19 | 25480 | 28302 | 2823 | forward |
| tail assembly chaperone | ERR2184201_19 | 25203 | 25463 | 261 | forward |
| tail protein | ERR2184201_14 | 24849 | 25478 | 630 | reverse |
| tail assembly chaperone | ERR2184201_19 | 24754 | 25122 | 369 | forward |
| BstA abortive infection system, replication inhibitor | ERR2184199_26 | 24273 | 25178 | 906 | forward |
| major tail protein | ERR2184201_19 | 24196 | 24708 | 513 | forward |
| tail terminator | ERR2184201_19 | 23790 | 24185 | 396 | forward |
| transcriptional repressor | ERR2184199_26 | 23489 | 24166 | 678 | forward |
| tail completion or Neck1 protein | ERR2184201_19 | 23455 | 23793 | 339 | forward |
| transcriptional repressor | ERR2184199_26 | 23182 | 23409 | 228 | reverse |
| minor head protein | ERR2184201_19 | 23145 | 23471 | 327 | forward |
| head-tail adaptor | ERR2184201_19 | 22819 | 23148 | 330 | forward |
| CDS | ERR2184199_26 | 22658 | 23038 | 381 | reverse |
| structural protein with Ig domain | ERR2184201_19 | 22544 | 22807 | 264 | forward |
| tail protein | ERR2184201_14 | 22513 | 24852 | 2340 | reverse |
| CDS | ERR2184199_26 | 22399 | 22566 | 168 | reverse |
| internal virion protein | ERR2184201_14 | 22053 | 22532 | 480 | reverse |
| internal virion protein | ERR2184201_14 | 21604 | 22068 | 465 | reverse |
| major head protein | ERR2184201_19 | 21603 | 22526 | 924 | forward |
| replication initiation O-like | ERR2184199_26 | 21312 | 22406 | 1095 | reverse |
| head scaffolding protein | ERR2184201_19 | 20988 | 21590 | 603 | forward |
| DnaB-like replicative helicase | ERR2184199_26 | 19936 | 21312 | 1377 | reverse |
| minor head protein | ERR2184201_19 | 19935 | 20861 | 927 | forward |
| CDS | ERR2184199_26 | 19713 | 19916 | 204 | reverse |
| CDS | ERR2184199_26 | 19475 | 19702 | 228 | reverse |
| CDS | ERR2184199_26 | 19258 | 19488 | 231 | reverse |
| endolysin | ERR2184201_14 | 19190 | 21604 | 2415 | reverse |
| CDS | ERR2184199_26 | 19034 | 19258 | 225 | reverse |
| unknown function CDS | ERR2184199_26 | 18581 | 19024 | 444 | reverse |
| portal protein | ERR2184201_19 | 18431 | 19930 | 1500 | forward |
| unknown function CDS | ERR2184199_26 | 18225 | 18584 | 360 | reverse |
| nucleotide kinase | ERR2184199_26 | 18004 | 18228 | 225 | reverse |
| CDS | ERR2184199_26 | 17864 | 18007 | 144 | reverse |
| unknown function CDS | ERR2184201_14 | 17454 | 19193 | 1740 | reverse |
| NinG/ Rap DNA junction specific endonuclease | ERR2184199_26 | 17299 | 17892 | 594 | reverse |
| CDS | ERR2184199_26 | 17136 | 17309 | 174 | reverse |
| terminase large subunit | ERR2184201_19 | 17117 | 18418 | 1302 | forward |
| CDS | ERR2184199_26 | 16955 | 17146 | 192 | reverse |
| terminase small subunit | ERR2184201_19 | 16672 | 17127 | 456 | forward |
| unknown function CDS | ERR2184201_19 | 16422 | 16646 | 225 | forward |
| anti-termination protein Q-like | ERR2184199_26 | 16344 | 16958 | 615 | reverse |
| holin/anti-holin | ERR2184199_26 | 15578 | 15895 | 318 | reverse |
| endolysin | ERR2184199_26 | 15181 | 15585 | 405 | reverse |
| transcriptional activator | ERR2184201_19 | 15154 | 15567 | 414 | forward |
| virion structural protein | ERR2184201_14 | 14872 | 17454 | 2583 | reverse |
| Rz-like spanin | ERR2184199_26 | 14783 | 15184 | 402 | reverse |
| unknown function CDS | ERR2184201_19 | 14612 | 14911 | 300 | forward |
| unknown function CDS | ERR2184201_19 | 14412 | 14615 | 204 | forward |
| unknown function CDS | ERR2184199_26 | 14200 | 14427 | 228 | reverse |
| unknown function CDS | ERR2184201_19 | 14175 | 14405 | 231 | forward |
| unknown function CDS | ERR2184201_19 | 13950 | 14144 | 195 | forward |
| unknown function CDS | ERR2184199_26 | 13777 | 14145 | 369 | reverse |
| CDS | ERR2184201_19 | 13735 | 13872 | 138 | forward |
| terminase small subunit | ERR2184199_26 | 13358 | 13780 | 423 | reverse |
| unknown function CDS | ERR2184201_19 | 13356 | 13754 | 399 | forward |
| unknown function CDS | ERR2184201_19 | 13078 | 13359 | 282 | forward |
| unknown function CDS | ERR2184201_19 | 12847 | 13065 | 219 | forward |
| unknown function CDS | ERR2184201_19 | 12533 | 12850 | 318 | forward |
| unknown function CDS | ERR2184201_14 | 12459 | 14807 | 2349 | reverse |
| unknown function CDS | ERR2184201_19 | 12228 | 12533 | 306 | forward |
| unknown function CDS | ERR2184201_19 | 12070 | 12231 | 162 | forward |
| terminase large subunit | ERR2184199_26 | 11944 | 13332 | 1389 | reverse |
| unknown function CDS | ERR2184201_19 | 11771 | 12073 | 303 | forward |
| unknown function CDS | ERR2184201_19 | 11614 | 11778 | 165 | forward |
| O-acetyltransferase | ERR2184201_14 | 11367 | 12470 | 1104 | forward |
| unknown function CDS | ERR2184201_19 | 11340 | 11609 | 270 | forward |
| DnaD-like helicase loader | ERR2184201_19 | 10516 | 11328 | 813 | forward |
| CDS | ERR2184201_19 | 10268 | 10498 | 231 | forward |
| integrase | ERR2184201_14 | 10008 | 11018 | 1011 | reverse |
| unknown function CDS | ERR2184201_14 | 9766 | 10011 | 246 | forward |
| portal protein | ERR2184199_26 | 9740 | 11941 | 2202 | reverse |
| unknown function CDS | ERR2184201_19 | 9582 | 10268 | 687 | forward |
| unknown function CDS | ERR2184201_14 | 9241 | 9663 | 423 | forward |
| Sak-like ssDNA annealing protein | ERR2184201_19 | 8905 | 9576 | 672 | forward |
| unknown function CDS | ERR2184201_14 | 8888 | 9244 | 357 | forward |
| head scaffolding protein | ERR2184199_26 | 8830 | 9729 | 900 | reverse |
| unknown function CDS | ERR2184201_19 | 8571 | 8912 | 342 | forward |
| unknown function CDS | ERR2184201_19 | 8387 | 8515 | 129 | forward |
| unknown function CDS | ERR2184201_14 | 8367 | 8891 | 525 | forward |
| CDS | ERR2184201_19 | 8245 | 8418 | 174 | forward |
| unknown function CDS | ERR2184201_19 | 8060 | 8248 | 189 | forward |
| unknown function CDS | ERR2184201_14 | 7963 | 8370 | 408 | forward |
| unknown function CDS | ERR2184201_14 | 7800 | 7970 | 171 | forward |
| HTH DNA binding protein | ERR2184201_19 | 7755 | 8039 | 285 | forward |
| major head protein | ERR2184199_26 | 7544 | 8815 | 1272 | reverse |
| excisionase | ERR2184201_19 | 7408 | 7707 | 300 | forward |
| unknown function CDS | ERR2184201_14 | 7312 | 7797 | 486 | forward |
| CDS | ERR2184201_87 | 7312 | 7437 | 126 | forward |
| unknown function CDS | ERR2184199_26 | 7299 | 7490 | 192 | reverse |
| DNA binding protein | ERR2184201_19 | 7190 | 7393 | 204 | forward |
| unknown function CDS | ERR2184201_14 | 7178 | 7327 | 150 | forward |
| unknown function CDS | ERR2184201_14 | 6846 | 7136 | 291 | forward |
| head-tail adaptor Ad3 | ERR2184199_26 | 6836 | 7318 | 483 | reverse |
| unknown function CDS | ERR2184199_65 | 6787 | 7026 | 240 | reverse |
| tail fiber assembly | ERR2184201_87 | 6740 | 7291 | 552 | forward |
| CDS | ERR2184199_65 | 6583 | 6747 | 165 | reverse |
| unknown function CDS | ERR2184201_14 | 6571 | 6774 | 204 | forward |
| anti-repressor Ant | ERR2184201_19 | 6429 | 7175 | 747 | forward |
| CDS | ERR2184199_65 | 6266 | 6547 | 282 | reverse |
| Lar-like restriction alleviation protein | ERR2184201_14 | 6149 | 6574 | 426 | forward |
| unknown function CDS | ERR2184201_19 | 6102 | 6353 | 252 | reverse |
| CDS | ERR2184199_65 | 5967 | 6269 | 303 | reverse |
| unknown function CDS | ERR2184201_19 | 5966 | 6121 | 156 | forward |
| CDS | ERR2184201_96 | 5960 | 6427 | 468 | reverse |
| unknown function CDS | ERR2184201_19 | 5727 | 5915 | 189 | reverse |
| unknown function CDS | ERR2184199_65 | 5674 | 5967 | 294 | reverse |
| unknown function CDS | ERR2184201_19 | 5572 | 5730 | 159 | forward |
| unknown function CDS | ERR2184199_65 | 5328 | 5681 | 354 | reverse |
| unknown function CDS | ERR2184199_65 | 5166 | 5321 | 156 | reverse |
| head closure Hc3 | ERR2184199_26 | 5083 | 6864 | 1782 | reverse |
| transcriptional regulator | ERR2184201_19 | 4952 | 5272 | 321 | reverse |
| unknown function CDS | ERR2184201_14 | 4909 | 6156 | 1248 | forward |
| unknown function CDS | ERR2184199_65 | 4804 | 5169 | 366 | reverse |
| unknown function CDS | ERR2184201_14 | 4742 | 4912 | 171 | forward |
| unknown function CDS | ERR2184201_96 | 4602 | 5360 | 759 | reverse |
| unknown function CDS | ERR2184201_14 | 4567 | 4770 | 204 | forward |
| metallo-protease | ERR2184201_19 | 4512 | 4934 | 423 | reverse |
| unknown function CDS | ERR2184199_65 | 4459 | 4800 | 342 | reverse |
| CDS | ERR2184201_96 | 4433 | 4600 | 168 | reverse |
| unknown function CDS | ERR2184201_14 | 4390 | 4548 | 159 | forward |
| tail needle protein | ERR2184199_26 | 4385 | 5083 | 699 | reverse |
| unknown function CDS | ERR2184201_19 | 4119 | 4424 | 306 | reverse |
| transcriptional regulator | ERR2184201_14 | 4044 | 4379 | 336 | forward |
| head morphogenesis | ERR2184199_26 | 3924 | 4385 | 462 | reverse |
| DNA methyltransferase | ERR2184199_65 | 3870 | 4316 | 447 | reverse |
| CDS | ERR2184201_96 | 3845 | 4336 | 492 | reverse |
| CDS | ERR2184201_178 | 3753 | 3830 | 78 | reverse |
| DksA-like zinc-finger protein | ERR2184199_65 | 3668 | 3862 | 195 | reverse |
| Mu Gam-like end protection | ERR2184201_14 | 3539 | 4036 | 498 | forward |
| long tail fiber protein distal subunit | ERR2184201_87 | 3485 | 6712 | 3228 | forward |
| DNA ejection | ERR2184199_26 | 3271 | 3930 | 660 | reverse |
| integrase | ERR2184201_19 | 2872 | 4011 | 1140 | reverse |
| hinge connector of long tail fiberprotein distal connector | ERR2184201_87 | 2811 | 3476 | 666 | forward |
| exonuclease | ERR2184201_14 | 2796 | 3542 | 747 | forward |
| CDS | ERR2184201_96 | 2774 | 3760 | 987 | reverse |
| DNA polymerase processivity factor | ERR2184201_178 | 2737 | 3747 | 1011 | reverse |
| host range and adsorption protein | ERR2184201_96 | 2529 | 2768 | 240 | reverse |
| CDS | ERR2184201_19 | 2346 | 2576 | 231 | reverse |
| unknown function CDS | ERR2184201_257 | 2294 | 2725 | 432 | forward |
| integrase | ERR2184199_65 | 2277 | 3434 | 1158 | reverse |
| unknown function CDS | ERR2184201_14 | 2179 | 2754 | 576 | forward |
| CDS | ERR2184201_297 | 2164 | 2367 | 204 | forward |
| CDS | ERR2184201_96 | 2084 | 2290 | 207 | reverse |
| unknown function CDS | ERR2184201_19 | 2041 | 2340 | 300 | reverse |
| CDS | ERR2184201_305 | 2029 | 2325 | 297 | forward |
| released from the phage upon host infection | ERR2184199_26 | 1912 | 3261 | 1350 | reverse |
| unknown function CDS | ERR2184201_14 | 1895 | 2032 | 138 | forward |
| DNA polymerase III theta subunit | ERR2184199_65 | 1857 | 2033 | 177 | reverse |
| unknown function CDS | ERR2184201_305 | 1795 | 2010 | 216 | forward |
| unknown function CDS | ERR2184201_14 | 1785 | 1898 | 114 | forward |
| long tail fiber protein proximal connector | ERR2184201_87 | 1633 | 2748 | 1116 | forward |
| CDS | ERR2184201_297 | 1613 | 2167 | 555 | forward |
| unknown function CDS | ERR2184201_14 | 1501 | 1788 | 288 | forward |
| unknown function CDS | ERR2184201_297 | 1398 | 1613 | 216 | forward |
| CDS | ERR2184201_297 | 1129 | 1401 | 273 | forward |
| unknown function CDS | ERR2184201_14 | 914 | 1504 | 591 | forward |
| DNA polymerase | ERR2184201_178 | 824 | 2737 | 1914 | reverse |
| tail protein | ERR2184201_257 | 728 | 2293 | 1566 | forward |
| integrase | ERR2184199_402 | 641 | 1156 | 516 | forward |
| CDS | ERR2184201_297 | 641 | 1132 | 492 | forward |
| unknown function CDS | ERR2184201_14 | 546 | 917 | 372 | forward |
| CDS | ERR2184201_178 | 428 | 619 | 192 | reverse |
| CDS | ERR2184199_402 | 336 | 644 | 309 | forward |
| unknown function CDS | ERR2184201_14 | 325 | 549 | 225 | forward |
| CDS | ERR2184201_19 | 305 | 1351 | 1047 | forward |
| integrase | ERR2184201_96 | 196 | 1554 | 1359 | reverse |
| Integrase H2C2 | ERR2184201_305 | 156 | 1805 | 1650 | forward |
| CDS | ERR2184201_178 | 143 | 418 | 276 | reverse |
| CDS | ERR2184201_96 | 101 | 199 | 99 | forward |
| CDS | ERR2184201_297 | 82 | 438 | 357 | forward |
| DNA methyltransferase | ERR2184199_402 | 2 | 301 | 300 | forward |
| DNA transfer protein | ERR2184199_26 | 2 | 1912 | 1911 | reverse |
| tail fiber protein proximal subunit | ERR2184201_87 | 2 | 1624 | 1623 | forward |
| CDS | ERR2184199_65 | 1 | 1809 | 1809 | forward |
| CDS | ERR2184201_19 | 1 | 312 | 312 | forward |
| tail protein | ERR2184201_257 | 1 | 750 | 750 | forward |

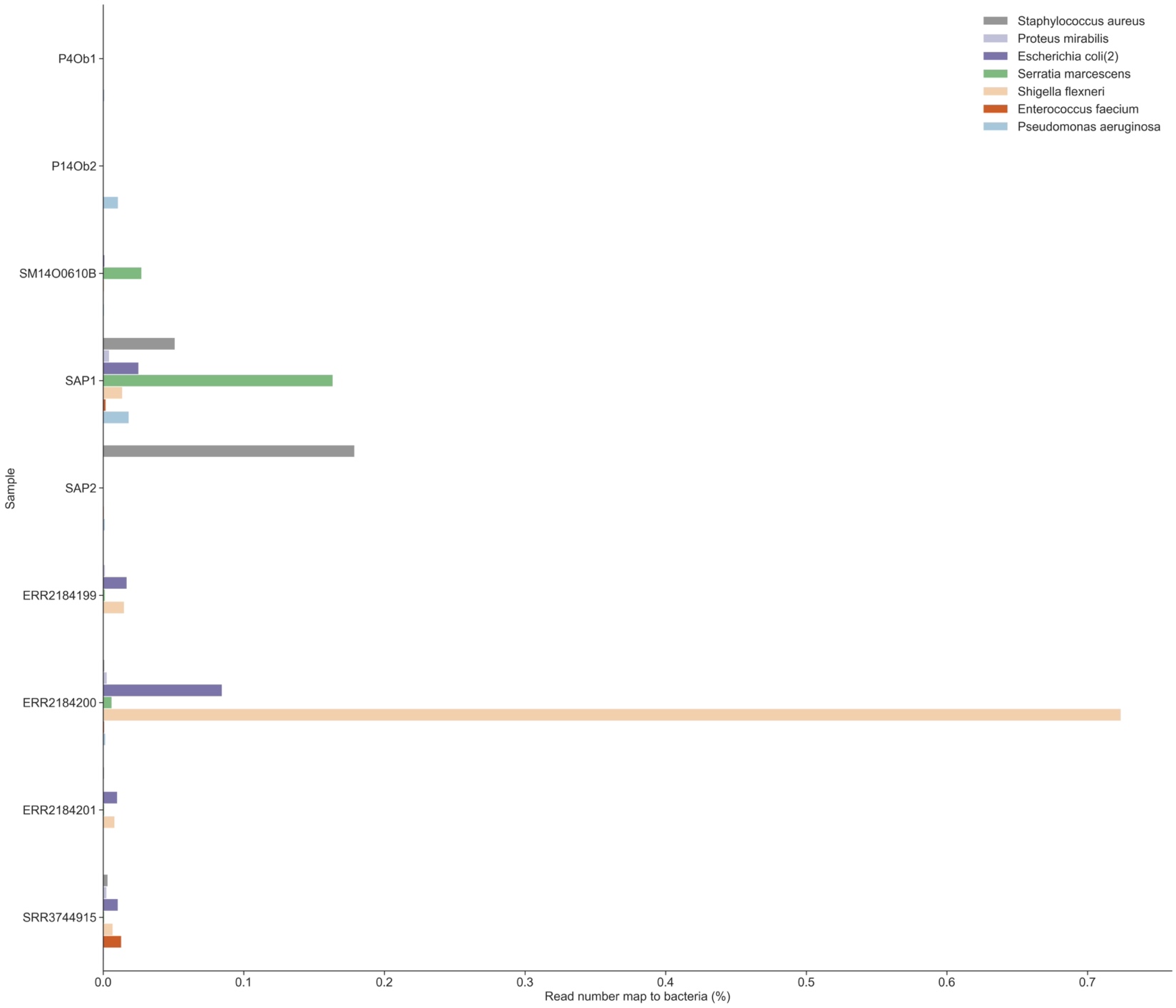

**Figure S1** Bacterial contamination in cocktails and individually sequenced phages

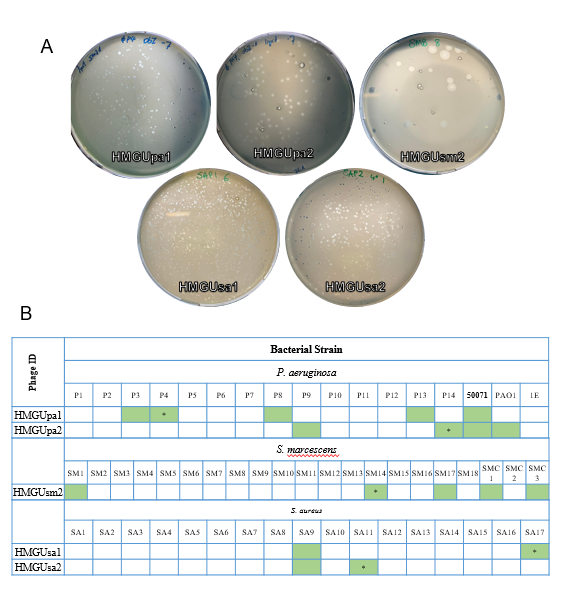

**Figure S2 A)** Plaque morphology of the five individually sequenced phages. **B)** Host range of the individually sequenced phages. Information about bacterial strains used for testing can be found in Table S4.

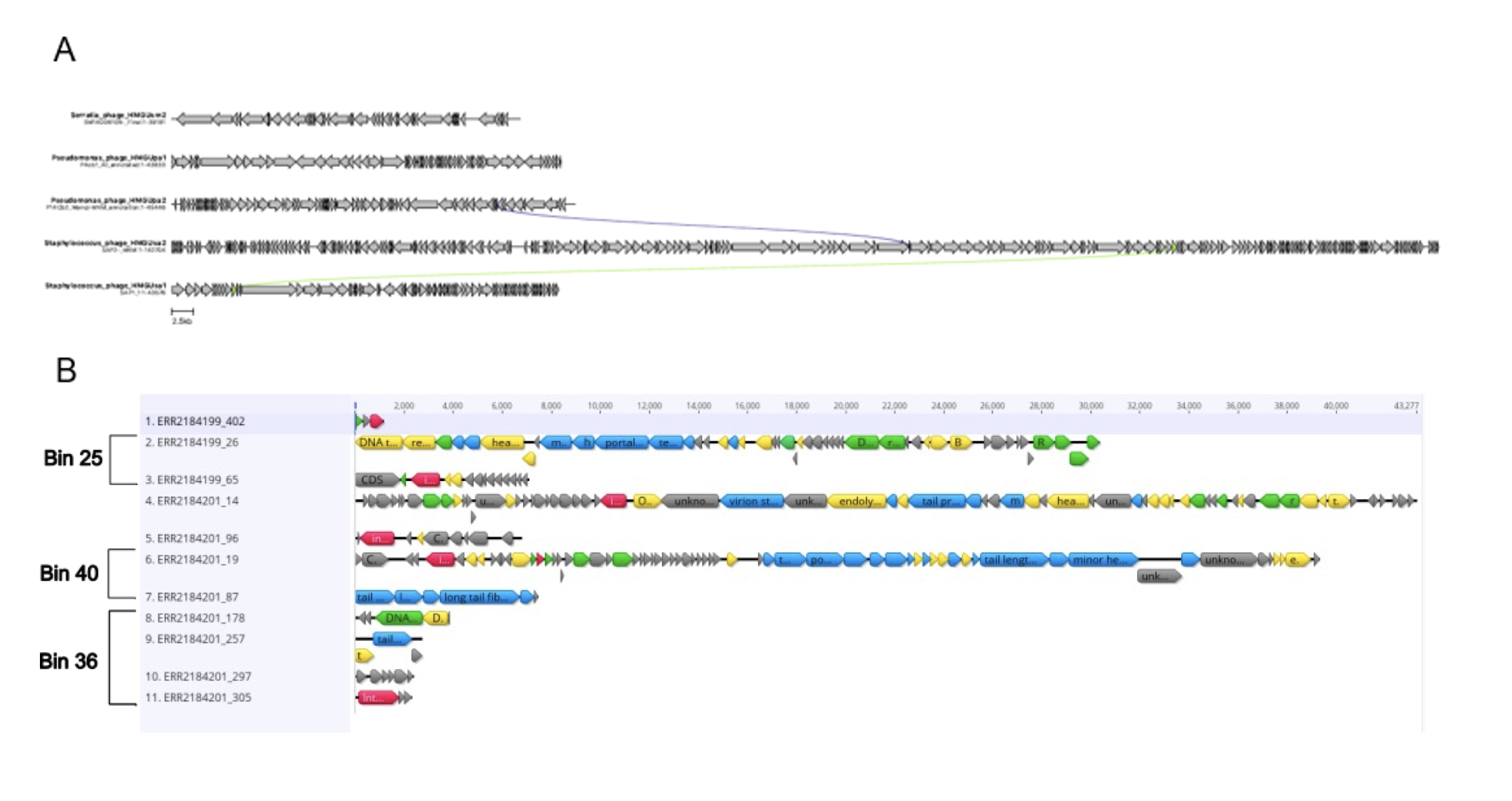
**Figure S3 A)** comparative genomic map of individually sequenced phages. **B)** Genomic map of four temperate contigs identified in the cocktails. Red denotes lysogeny related genes, blue shows structural genes, and green shows genes involved in nucleotide metabolism. Genes with an unknown function are shown in grey and other genes are shown in yellow.
